# Multiplicity of *Agrobacterium* infection of *Nicotiana benthamiana* for transient DNA delivery

**DOI:** 10.1101/2023.02.07.527564

**Authors:** Erik D. Carlson, Jakub Rajniak, Elizabeth S. Sattely

**Affiliations:** Department of Chemical Engineering, Stanford University, Stanford, California 94305, USA; Department of Bioengineering, Stanford University, Stanford, California 94305, USA; Howard Hughes Medical Institute, Stanford University, Stanford, California 94305, USA

## Abstract

Biological DNA transfer into plant cells mediated by *Agrobacterium* represents one of the most powerful tools for the engineering and study of plant systems. Transient expression of transfer DNA (T-DNA) in particular enables rapid testing of gene products and has recently been harnessed for facile combinatorial expression of multiple genes. In analogous mammalian cell-based gene expression systems, a clear sense of the multiplicity of infection (MOI) allows users to predict and control viral transfection frequencies for applications requiring single vs. multiple transfection events per cell. Despite the value of *Agrobacterium*-mediated transient transformation of plants, MOI has not been quantified. Establishing MOI for *Agrobacterium* T-DNA delivery at the level of single plant cells would allow users to design genomic library delivery conditions (at most 1 event/cell), or maximize co-delivery of T-DNA loads from separate *Agrobacterium* (>1 event/cell). Here, we analyze the Poisson probability distribution of T-DNA transfer in leaf pavement cells to determine the MOI for the widely used model system *Agrobacterium* GV3101/*Nicotiana benthamiana*. These data delineate the relationship between an individual *Agrobacterium* strain infiltration OD_600_, plant cell perimeter and leaf age, as well as plant cell co-infection rates. Our analysis also establishes experimental regimes where the probability of near-simultaneous delivery of >20 unique T-DNAs to a given plant cell remains high throughout the leaf. We anticipate that these data will enable users to develop new approaches to in-leaf library development using *Agrobacterium* transient expression and the reliable combinatorial assaying of multiple heterologous proteins in a single plant cell.

## Introduction

Stable expression of multiple transgenes is a powerful way to endow a plant with a new trait such as resistance to potato blight ^1^ or the production of polyunsaturated fatty acids in soy ^2^. These strategies require the coordinated action of multiple transgenes. However, many challenges exist for multigene expression systems, including a lack of suitable genetic parts (promoters, terminators, etc.), rapid methods for integrating multiple genes, and lengthy design-build-test cycles required for optimizing pathways and gene sets.

*Agrobacterium*-mediated DNA delivery is one of the most widely used tools in plant biotechnology. *Agrobacterium* transfers single-stranded DNA into plant cells as part of the infection process; this DNA is ultimately integrated into the host genome for sustained expression, termed “stable expression”. This process has been harnessed for the delivery of user-defined cargo, and has become a major method for generating transgenic plant lines when DNA is delivered into undifferentiated tissue ^3^. While numerous species across the plant kingdom are known to be hosts for *Agrobacterium* infection, some are amenable to high infection rates with low symptom development. In a few of these plants, DNA can be expressed transiently in somatic plant tissues by infiltrating a suspension of bacterial cells into leaf tissue (Figure 1A).

**Figure 1.**
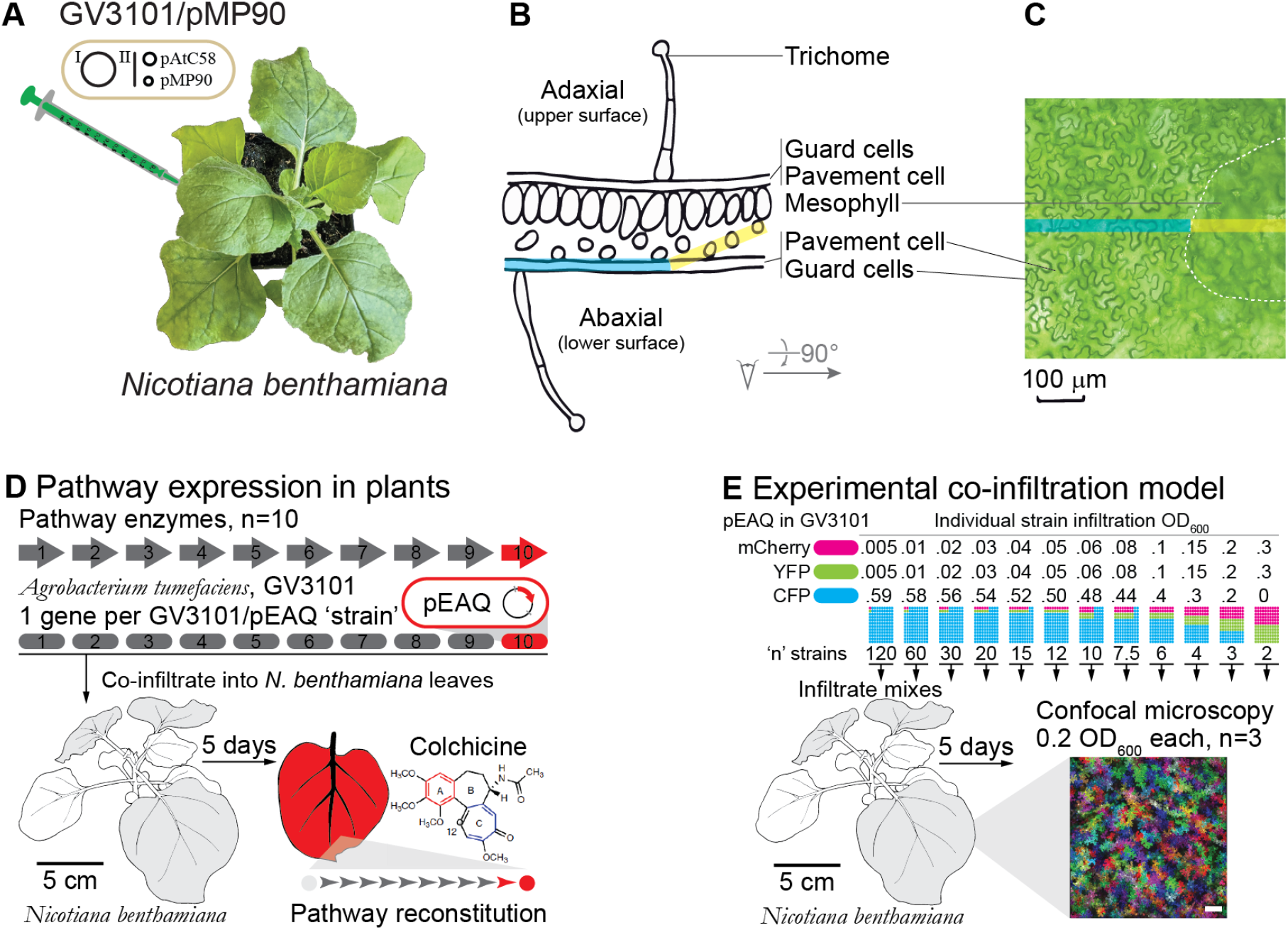
Transient expression of multiple genes in *N. benthamiana* leaves by co-infiltration of *Agrobacterium* expression strains. **A**. Infiltration of *Agrobacterium tumefaciens* strain GV3101 into *Nicotiana benthamiana* leaves **B**. Cross-section diagram of *N. benthamiana* leaf based on microscope images of typical leaves used in this experimental system. **C**. Brightfield image of abaxial (lower surface) side showing pavement cells where data is collected for this study. **D**. Application of transient expression for reconstitution of multi-step metabolic pathways in *N. benthamiana* leaves. Each gene is delivered in a separate *Agrobacterium* strain; infiltration of strain mixtures into leaves results in heterologous protein expression and metabolite production^9^. **E**. Strategy to quantify the number of unique *Agrobacterium*-derived T-DNA gene products that are present in a given plant cell. Genes for three orthogonal fluorescent proteins are each carried by the T-DNA in the pEAQ plasmid of individual strains. Strain mixtures are co-infiltrated into leaves at various ratios keeping total infiltration OD_600_ at 0.6. Confocal microscopy allows identification of co-infection events at a single cell level. Confocal microscopy image of 0.2 OD_600_ per strain, n=3. White scale bar is 100 μm.

This strategy has been widely used for rapidly testing transgene function and even as a method for heterologous protein production without the need for generating stable transgenic plant lines^4^. A particularly effective pairing is *Agrobacterium* strain GV3101(pMP90)^5^ with binary vector pEAQ ^6^ in the plant host *Nicotiana benthamiana* (Figure 1A, Supplementary Figure 1A,B). *N. benthamiana* grows relatively quickly, with ample leaf material for easy infiltration in ~4-6 weeks after planting. GV3101/pEAQ infiltration into *N. benthamiana* leaves leads to detectable T-DNA product expression in a matter of days and heterologous protein levels that have been reported to reach upwards of 1.5 g kg^−1^ fresh weight (FW) of GFP, and 325 mg kg^−1^ FW of human antibody 2G12 ^6^.

Notably, it has been found that co-infiltration of multiple *Agrobacterium* strains with unique expression constructs leads to simultaneous expression of the T-DNAs in plant leaves^7^. This has enabled the rapid testing of multigene biosynthetic pathways ^8–11^. The modularity of this co-infiltration method, and the fact that each gene can be driven by the same promoter (e.g. 35S) often makes this approach for pathway reconstitution in *N. benthamiana* leaves more straightforward compared to traditional heterologous hosts such as *E. coli* and yeast, which also have limits with the complexity of heterologous proteins produced compared to a plant system. Furthermore, this type of rapid, combinatorial transient expression in plants has drastically accelerated the design-build-test cycle for testing plant pathways prior to the generation of stable transgenic lines that can require months to years of development.

Multi-step pathways (>10 unique gene products required, Figure 1D) are expressed routinely, and accumulation of the expected products is often observed. For example, co-infiltration of 16 *Agrobacterium* GV3101(pMP90)/pEAQ strains delivering a 16 gene pathway transiently in *N. benthamiana* leaves produces (—)-deoxypodophyllotoxin up to 4.3 mg g^−1^ dry weight^10^. Notably, intermediates that would be the result of “partial” pathways are typically not observed. This may be due to enzyme specificity (cells with fewer than all required genes do not make new metabolic products), metabolite sharing across cells, and/or high efficiency of T-DNA delivery with enough cells receiving a complete set of unique gene constructs.

In analogous experimental strategies for rapid genetic manipulation of cells, such as viral transfection of mammalian cells in culture, multiplicity of infection (MOI) can be a critical experimental design parameter to help predict how many unique transfection events a given target cell is likely to undergo. MOI was developed with bacteriophage/bacteria systems^12^ and is now routinely used in viral/mammalian transformation systems ^13^. Despite the importance of *Agrobacterium*-mediated transient expression as a widely used tool to investigate plant cell biology and pilot transgene constructs, the process by which multiple T-DNAs function in a coordinated way is not well understood and no analogous MOI value has been determined for this system. For biosynthetic pathway reconstitution using this approach, the observed robust pathway product formation by co-infiltration of multiple *Agrobacterium* strains in *N. benthamiana* leaves suggests high rates of co-infection at the plant cell level and/or high rates of metabolite or nucleic acid sharing across plant cells, but this has not been extensively studied.

Here we develop an *in planta* co-infiltration fluorescence assay to quantify co-infection rates (Figure 1E) and a statistical model that allows us to predict the number of unique T-DNA products present in each plant pavement cell, an MOI metric for this system. We predict that a typical *N. benthamiana* plant pavement cell in the top three fully expanded leaves can receive on average 1-3 infection events by a given *Agrobacterium* strain infiltrated at 0.2 OD_600_. Our work suggests that a high rate of co-infection by co-infiltrated strains is a main driver for observed pathway reconstruction in this transient expression system. These data also inform the design of an experimental design regime where at most 1 T-DNA type enters a given cell. This problem is analogous to MOI measurements for bacteriophage/bacteria and virus/mammalian cell systems that are commonly used for genetic screens. This is an essential first step towards the development of novel genetic screens in whole plant leaves, where at most 1 T-DNA type/plant cell would be needed. More broadly, we anticipate that quantification and modeling of multi-gene delivery through *Agrobacterium* transient expression can contribute to accelerating the design-build-test cycle for engineering plant traits.

## Results

### GV3101(pMP90)/pEAQ strain tracking and T-DNA expression timing

To capture early stages of infection and get a sense of timing for fluorescent protein expression, we first tracked *Agrobacterium* simultaneously with plant-cell expressed T-DNA after leaf infiltration, as in 14. We constructed a version of pEAQ-GFP (T-DNA containing green fluorescent protein) with an added constitutive red fluorescent protein (RFP) expression cassette ^15^ on the non-T-DNA backbone, which turns the colony and cell pellet visibly pink (Figure 2A). This fluorescently tags *Agrobacterium* that maintains the pEAQ plasmid with RFP, and GFP encoded in the T-DNA is only present once T-DNA is delivered to and expressed by a plant cell. This strategy enables us to track the strain during the infection time course and expression of T-DNA products by the infected plant cell.

**Figure 2.**
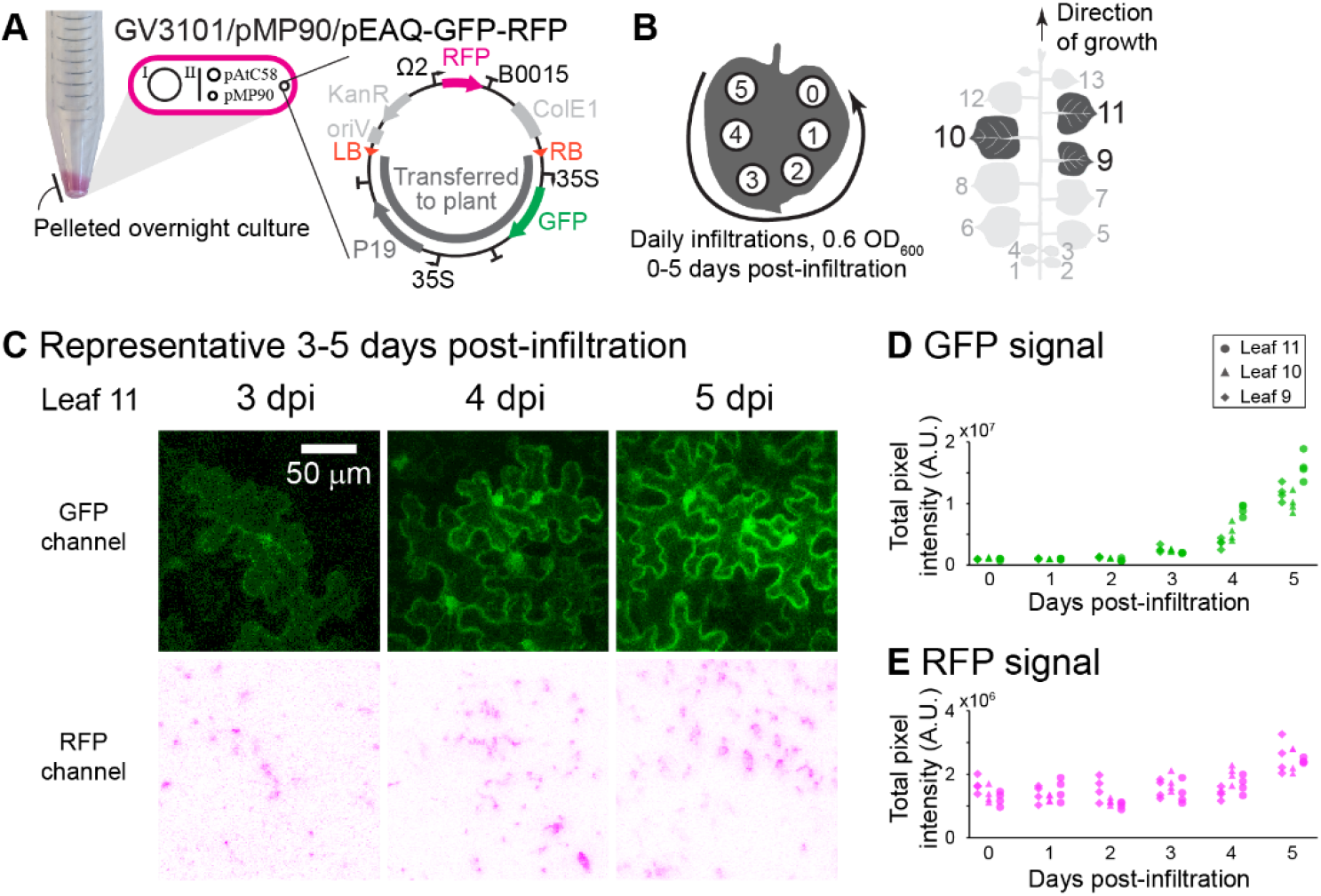
Fluorescence microscopy of RFP-expressing *Agrobacterium* with GFP T-DNA load in *N. benthamiana* leaves. **A**. A constitutive RFP cassette was added to the non-T-DNA portion of a pEAQ vector with GFP T-DNA. **B**. Acetosyringone-induced bacterial cultures were infiltrated daily into different spots on the same leaves. **C**. Representative 3, 4 and 5 days post-infiltration fluorescence microscopy images of leaf 11. The GFP channel shows characteristic cytosolic expression in the plant pavement cells, and the RFP channel marks *Agrobacterium* location in the leaf. RFP channel images are shown as inverse negatives of a green false colored image, to display magenta on a white background for visual clarity. **D**. GFP channel pixel intensity. **E**. RFP channel pixel intensity counts. Each data point corresponds to the corresponding channel from a single image.

Once a day for 5 days and once on the morning of imaging, 0.6 OD_600_ acetosyringone-induced GV3101(pMP90)/pEAQ-GFP-RFP was infiltrated into different sites on the same top 3 fully-expanded *N. benthamiana* leaves 9, 10 & 11 (Figure 2A,B, Supplementary Figure 4A,B). 5 days after the first infiltration, sites were imaged on the underside of the leaf using a Leica SP8 confocal microscope. Laser powers were set for the 5 days post infiltration (dpi) spot on leaf 11 and kept constant for imaging all the remaining spots (Supplementary Figure 4C,D). Representative images of the 3, 4 and 5 days post-infiltration spots show RFP-expressing *Agrobacterium* throughout the image with detectable GFP expression in the plant pavement cells starting at 3 dpi (Figure 2C, Supplementary Figure 4C). The GFP channel is characteristic of cytosolic expression showing a clear cell outline and nucleus, caused by the vacuole taking up the majority of the cell volume thus pushing the cytosol to the cell edges and around the nucleus. Quantifying pixel intensity counts for GFP (Figure 2D) shows protein expression can be detected at 3 days post-infiltration, with a marked increase in fluorescent signal at 5 dpi. *Agrobacterium* RFP signal is observed *in planta* in all infiltration conditions (Supplementary Figure 4D), with an apparent increase at 5 dpi concomitant with protein expression (Figure 2E). We chose 5 dpi for co-infiltration experiments given the clear signal to noise ratio, the early stage of infection, the lack of an apparent defense phenotype or tissue damage in the leaves, and the fact that biosynthetic pathway reconstitution experiments are typically performed in this timeframe.

### Determination of Agrobacterium infection frequencies and its relationship to infiltration OD_600_

Multiplicity of infection is classically defined as the ratio of infectious agents to infection targets, termed MOI_input_. The more accurate and useful metric, MOI_actual_, is derived from experimental data and thus accounts for dynamics of vector adsorption to the target cell and subsequent infection success. For the *Agrobacterium*-mediated transient expression system, we define an MOI_actual_ metric based on the relationship of infiltration OD_600_ to co-infection frequencies. Specifically, we use a Poisson distribution model with parameter λ defined as the product of strain infiltration OD_600_ (*A_x_*), and a fitted constant term α, defined as the mean number of unique T-DNA products delivered by an *Agrobacterium* strain to a given plant cell per infiltration OD_600_ for a given plant pavement cell (Equations 1 & 2, Poisson Distribution modeling experimental section). Accordingly, we anticipate that the MOI should change depending on the total number of *Agrobacterium* cells infiltrated into a leaf.

To examine and quantify the transfer of unique T-DNAs in this type of combinatorial, co-infection process, we chose to use *Agrobacterium* strains each encoding a different fluorescent protein with an orthogonal emission signal distinguishable by confocal microscopy (Figure 3A, Supplementary Figure 1C). Specifically, cyan, yellow, and red fluorescent proteins (CFP, YFP, mCherry respectively, optimized for cytosolic expression in plant cells ^16^) were cloned into the pEAQ-HT vector ^6^ under the 35S promoter, and transformed into *Agrobacterium tumefaciens* GV3101(pMP90). Imaging by confocal microscopy then allows us to detect and enumerate pavement cells that contain different combinations of fluorescent protein products after a specified time interval. To determine α, the infection frequency as a function of *Agrobacterium* concentration, we varied inoculation OD_600_ and tracked expression of T-DNA fluorescent protein products. We noted that most, but not all pavement cells express fluorescent protein when leaves are infiltrated with a saturating OD_600_ of *Agrobacterium* (Supplementary Figure 9). We designated *Agrobacterium* containing either YFP- and mCherry-encoded T-DNA as “indicator” strains, and used CFP as a dilution strain representing a bulk population and as a marker for the total number of infectable plant pavement cells (Figure 3A,B). While we focused our analysis on pavement cells due to ease of imaging, we noted that mesophyll cells in plant leaves are also infectable (Supplementary Figure 3).

**Figure 3.**
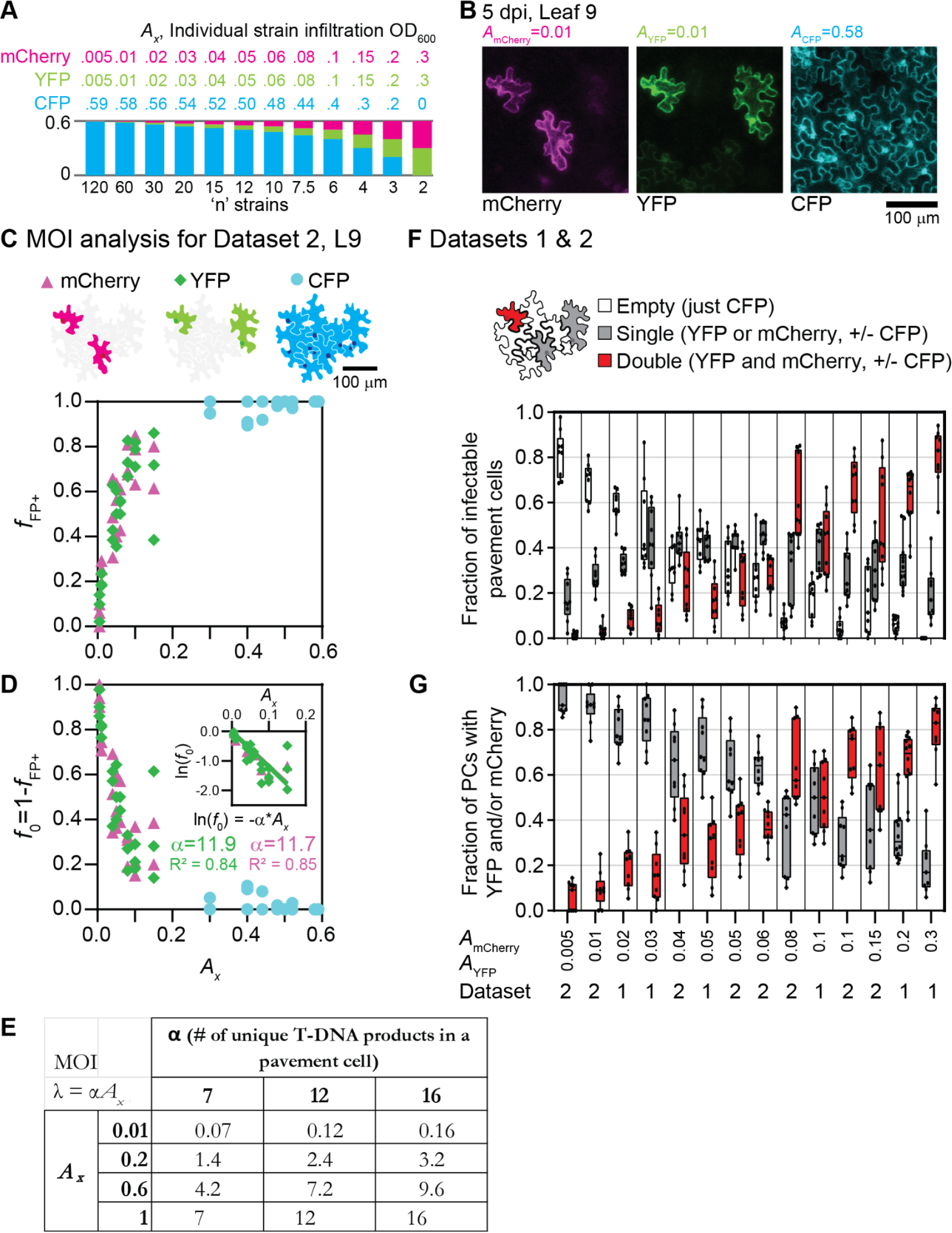
Quantification of infected cells at different OD_600_s allows for determination of MOI. **A**. One fluorescent stain (GV3101/pEAQ-CFP) used as a dilution strain, keeping total *Agrobacterium* infiltration OD_600_ constant at 0.6. *Agrobacterium* mixes were infiltrated into the top 3 fully expanded leaves of 6-week old *N. benthamiana* plants (leaves 8, 9 and 10), and imaged after 5 days. Minimum of 3 images were collected for each leaf, with minimum 9 images total for each co-infiltration condition. **B**. Selected image from Leaf 9, infiltration mix n = 60, Dataset 2. **C-E**. Multiplicity of infection analysis for Leaf 9 from Dataset 2. **C**. Fraction of infectable pavement cells positive for a given fluorescent protein (*f*_FP+_) dependent on strain infiltration OD_600_. **D**. Fraction of infectable pavement cells negative for a given fluorescent protein, *f*_0_=1-*f*_FP+_. Inset: fitting reduced Poisson distribution model (equation 5) to calculate α values for mCherry and YFP. **E**. Multiplicity of infection (λ) values for various infiltration OD_600_ and α values. **F**. Box and whisker plots show fraction of infected pavement cells that are empty (just CFP), single infected (just YFP or mCherry, with or without CFP), and double infected (YFP and mCherry, with or without CFP), at various indicator strain infiltration OD_600_, *A_x_*. **G**. Box and whisker plots for only YFP and/or mCherry positive plant pavement cells (PCs), single or double infected at various indicator strain infiltration OD_600_.

Individual strain OD_600_s were varied as illustrated in Figure 3A with the total infiltration OD_600_ kept constant at 0.6. Induced strain mixes were hand infiltrated into the top 3 fully expanded leaves of 6-week-old *N. benthamiana* (leaves 8, 9 and 10, counting up from the cotyledons as 1 and 2), and imaged at 5 dpi (Supplementary Figures 5C and 6A). At least 3 images were collected per leaf, with at least 9 images total for each co-infiltration condition. Images were manually analyzed using ImageJ, to quantify the total number of cells expressing each of the three fluorescent proteins as well as the combination of fluorescent protein signals in a given plant pavement cell (Supplementary Tables 1 and 2).

For the MOI calculation, we considered any cell that contained at least one fluorescent protein as “infectable” and counted the fraction of those cells that contained CFP, mCherry, and YFP (*f*_FP+_) as a function of each strain infection OD_600_ (Figure 3C). As described in the Poisson distribution modeling equations experimental section, the reduced Poisson distribution of X=0 can be applied to cells not expressing the fluorescent protein, *f*_0_ = 1-*f*_FP+_, (Figure 3D, Supplementary Figure 5D-F, Supplementary Figure 6B-D), to fit for α, and thus MOI_actual_. We found that MOI can vary from between ~0.1-16 for various individual strain OD_600_s (Figure 3E), revealing that at high infiltration OD_600_, 16 or more unique T-DNAs are expected to be delivered on average to a given plant cell. We also noted that the calculated α values changed depending on leaf age. Older leaves, e.g. L8 were found to have a higher α (and corresponding MOI) relative to L10 (Supplementary Figure 5D-F, Supplementary Figure 6B-D).

A few assumptions we are making are as follows: first, does fluorescent protein detection in a cell correspond to *Agrobacterium* infection and T-DNA transfer in that same cell? This is probably not always the case, as we know proteins of the size of RFP can travel from one plant cell to another via the plant’s plasmodesmata^17^. Independence of infection events, meaning infection of a plant cell by one *Agrobacterium* cell does not affect the likelihood of a second *Agrobacterium* cell binding and infecting the same plant cell when co-infiltrated, is supported by Chi-square analysis of independence of infection frequencies of the two indicator strains, pEAQ-YFP and pEAQ-mCherry (Supplementary Tables 1 and 2). Also of note, here we are not considering intensity of expression, which changes depending on the total number of T-DNA molecules delivered into a cell and the cell-to-cell variation in expression dynamics^18^. In this work, these data reflect only binary yes/no presence of the fluorescent protein in a given plant pavement cell which could result from one or more infection events with the same strain.

Filtering the data to focus on indicator strains YFP and mCherry reveals useful experimental design spaces (Figure 3F,G). Above individual strain infiltration OD_600_ of ~0.1, we noted that the majority of plant pavement cells are infected and positive for both YFP and mCherry. These data point to high rates of co-infection upon co-infiltration of multiple *Agrobacterium* strains encoding for pathway enzymes as a main driver of the successful biosynthetic pathway reconstitution, where our lab typically uses and infiltration OD_600_ of 0.2-0.3 per *Agrobacterium* strain. The data also helps inform use cases such as genetic library delivery where, when plant cells are infected, the delivery of at most one T-DNA type per cell is desired. For these experiments, infiltration OD_600_s of less than ~0.02 result in a majority of infected cells with only 1 of the indicator strains (Figure 3G). However, this is at a tradeoff with total number of plant cells infected under these conditions, e.g. ~20% of infectable pavement cells for *A_x_* = 0.005 contain YFP or mCherry (Figure 3F), with the majority of those cells with only one of the indicator strains (Figure 3G).

### High rates of co-infection correspond to increasing plant pavement cell perimeters

With a method established for quantifying co-infection with 3 individual strains, we next examined which major variables influenced MOI across the plant body, and specifically focused on leaf age. As before, we mixed and co-infiltrated three *Agrobacterium* strains, each bearing T-DNA for a different fluorescent protein but extended our analysis to include leaves 6-13. The 3 strains (encoding for CFP, YFP or mCherry) were mixed at equal volumes, for a final concentration of 0.2 OD_600_/strain, 0.6 OD_600_ total (Figure 4A and Supplementary Figure 8A, 1:1:1). Separately, we also installed the *Agrobacterium*-labeling constitutive RFP cassette into the pEAQ-YFP vector backbone, creating pEAQ-YFP-RFP. This strain was induced and mixed with the pEAQ-CFP strain to visualize *Agrobacterium* & T-DNA expression at infiltration OD_600_ 0.2 (Supplementary Figure 8A, 2:1). Imaging the underside of the leaf at 5 dpi, we observed plant pavement cells expressing the cytosolic fluorescent proteins from T-DNA loads (Figure 4B, Supplementary Figure 8C), and *Agrobacterium* tagged with RFP (Supplementary Figure 8D). Representative images from leaves 13 (youngest) and 6 (oldest) show robust expression of proteins from T-DNA constructs, and that the majority of plant pavement cells are infected (Figure 3B).

**Figure 4.**
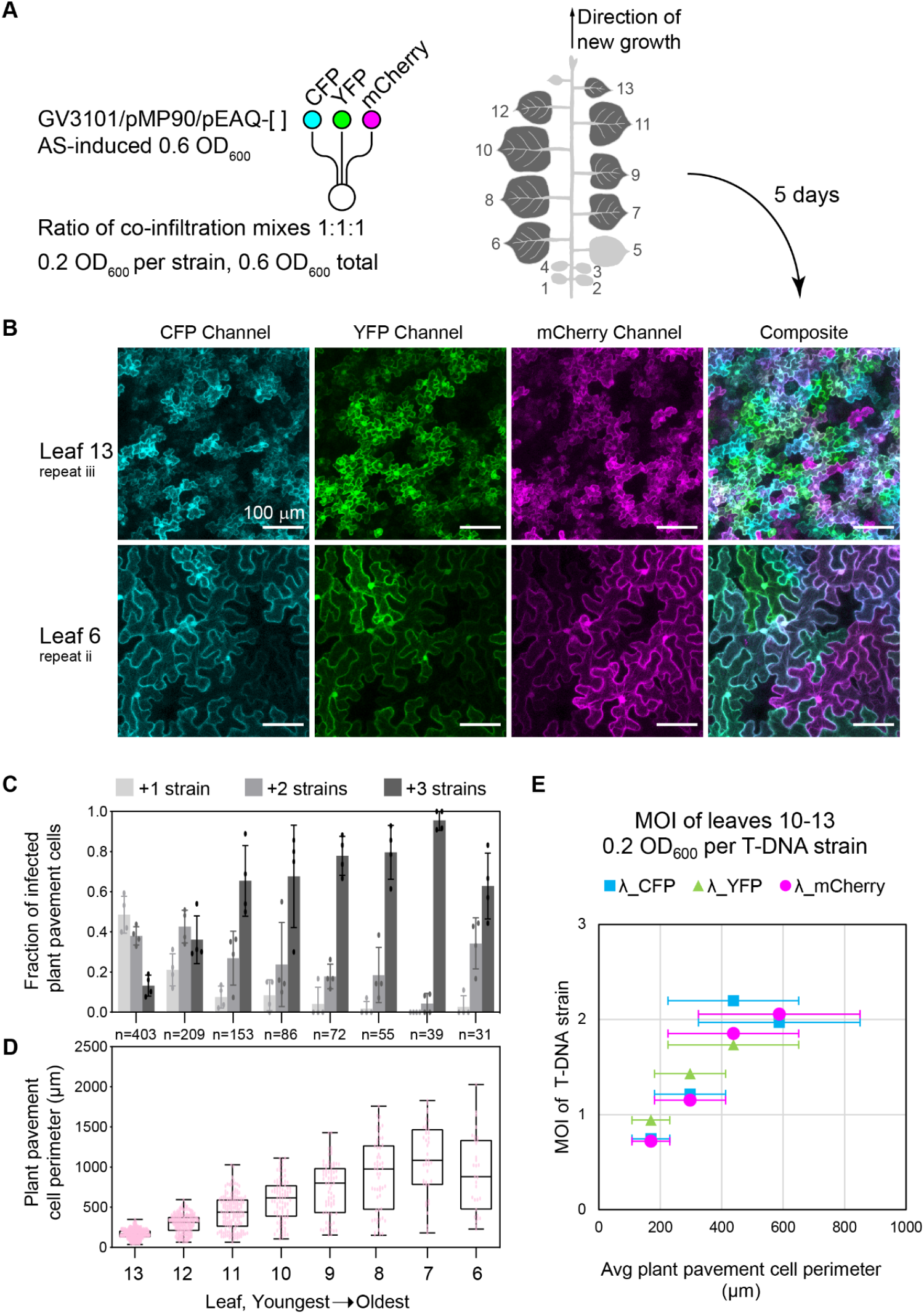
Quantification of pavement cell co-infection rates of *Agrobacterium* at equal ODs and the effect of plant cell size. **A**. Experimental setup and infiltration into leaves 6-13 of a 6 week old *Nicotiana benthamiana* plant. Three *Agrobacterium* strains, each containing a gene for a different fluorescent protein were co-infiltrated at equal OD_600_ (0.2 OD_600_/strain, 0.6 OD_600_ total, 1:1:1). 6-week old *Nicotiana benthamiana* plant structure, infiltrating leaves 6-13 as counted up from the cotyledons (1 & 2). **B**. Representative confocal microscopy images from leaves 13 and 6, of 1:1:1 co-infiltration condition of CFP:YFP:mCherry at 5 dpi. **C**. Fraction of infected plant pavement cells that contain one, two, or three fluorescent proteins. Averages are 4 images each from a different region of the same plant leaf disc +/− SD. **D**. Box and whisker plots of plant pavement cell perimeters (μm) by leaf. Each data point is the perimeter of an individual plant pavement cell. Number of plant pavement cells fully in the 4 image frames per condition is noted by “n”. **E**. Average pavement cell perimeters vs calculated multiplicity of infections for CFP, YFP and mCherry. Error bars show +/− SD.

In each microscopy image for the 1:1:1 co-infiltration case, the perimeter and number of T-DNA products present was quantified for plant pavement cells fully in the microscope image (Supplementary Figures 9 and 10A). Infected plant pavement cells were binned as single, double and triple infected based on the total different fluorescent proteins observed in that cell (Figure 4C). In leaves 6-13, the majority of plant pavement csells are positive with at least 1 fluorescent protein (Supplementary Figures 8C and 10A). For older leaves (e.g. leaves 6-7), we observed a majority of infected cells with all 3 fluorescent proteins (Figure 4C) that appeared to track with the increased perimeter of these pavement cells (Figure 4D). Indeed, quantifying MOI, we observed good correlation with average plant cell perimeter (Figure 4E). Data for cells >600 um was not included for two (interrelated) reasons: (1) There are too few cells of this size that fit in a single image to give an accurate estimate, and (2) essentially all cells are triply infected in the older leaves, which also precludes getting an accurate estimate of MOI. Together, these data suggest a simple statistical model where the increase in cell perimeter increases the likelihood of *Agrobacterium* infection. Our results also highlight the importance of comparing leaves of comparable developmental stage for applications that rely on successful combinatorial expression of multiple distinct T-DNAs (e.g. biosynthetic pathway reconstitution).

### Exploring the theoretical co-infiltration design space

Based on the probability model determined by fitting the data to a Poisson distribution, we next explored possible experimental regimes for co-infection by n > 2 co-infiltrated strains. Towards this end, we generated plots based on the Poisson distribution analysis that represent likely experimental design for the GV3101(pMP90)/pEAQ in *N. benthamiana* system.

Figure 5A plots various Poisson distribution curves (equation 1), with overlaid observed co-infection data for indicator strains YFP and mCherry, showing good agreement between the model and our observed data (Figure 5A, Supplementary Figure 11). Figure 5B shows the number of co-infiltrated strains n related to the probability of a given plant pavement cell being infected by all ‘n’ strains. Curves of various total infiltration OD_600_ representing typical co-infiltration experimental design spaces are shown (*A_Total_*= 0.6, 1.6, 4 and 10 OD_600_), calculated with α = 7, 12 and 16 mean number of infection events in a given plant pavement cell per strain infiltration OD_600_. Reflective of typical co-infiltration experiments, noted on the graph are cases of individual strain infiltration OD_600_ of 0.2 for each total infiltration OD_600_ curve. The black bars extending up and down from the circled point show the possible range of complete co-infection by all n strains, depending on the MOI_actual_ value for that given experiment. These data are in line with observed yields of 4.3 mg g^−1^ dry weight of a target medicinal compound, produced by co-infiltration of 16 unique *Agrobacterium* strains into *N. benthamiana* leaves at a total OD_600_ of 3^10^; our model predicts that the probability that T-DNAs expressed in a given cell approaches 50%, especially for infiltrations of young leaves.

**Figure 5.**
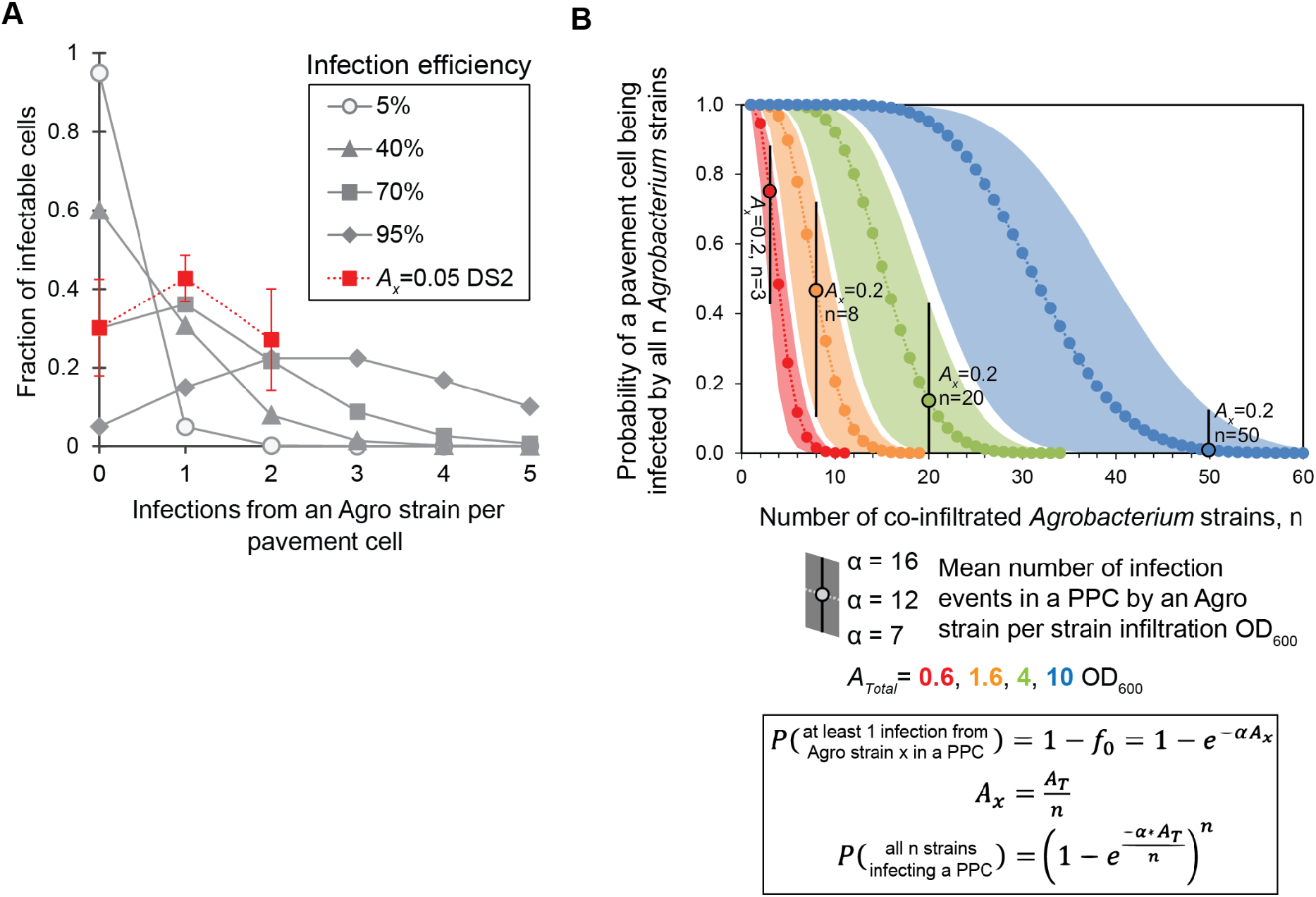
Poisson distribution modeling to estimate multiplicity of infection for GV3101 in *N. benthamiana*. **A**. Expected number of infection events per plant pavement cell at various infection efficiencies based on Poisson distribution (equation 1). Curves with solid connection lines are based on the Poisson distribution form, with overlaid comparison of experimental data in red square with dashed line. **B**. Probability of a plant pavement cell being co-infected by all n co-infiltrated *Agrobacterium* strains, fixed total infiltration OD_600_, *A_Total_* of 0.6, 1.6, 4, and 10. Representative range of α values used to generate curves. Points circled in black (α=12) with black bars extending to upper and lower α values note the co-infiltration conditions with individual strain OD_600_ of 0.2 for a given total infiltration OD_600_.

## Discussion

*Agrobacterium*-mediated transient expression in *Nicotiana benthamiana* is a powerful tool for combinatorial expression and discovery of heterologous pathway enzymes. Our data point to high rates of co-infection by co-infiltrated *Agrobacterium* strains to be a main driver of the platform’s utility in this space. By modulating the number and infiltration OD_600_ of *Agrobacterium* carrying different T-DNA loads, we show an ability to control key infection metrics like co-infection of a plant cell by different T-DNA loads. We have a clearer sense of the accessible experimental landscape of this GV3101(pMP90)/pEAQ/*N. benthamiana* system for designing co-infiltration experiments.

Our datasets focus on plant pavement cells, due to the data collection limitations of confocal microscopy slide preparation and the number of infiltration conditions we targeted. The bulk of a *Nicotiana benthamiana* leaf tissue are mesophyll cells that do get infected upon GV3101/pEAQ infiltration (Supplementary Figure 3) and are likely where most of the heterologous expression occurs. Generally, mesophyll cells are bigger than plant pavement cells and have more exposed surface area for *Agrobacterium* binding, so we postulate that co-infection rates would generally be higher in mesophyll cells compared to pavement cells. Therefore, MOI based on plant pavement cells would likely be an under-estimation of MOI in mesophyll cells.

Towards understanding the dynamics of heterologous pathway reconstitution in *N. benthamiana*, our statistical modeling predicts that co-infiltration of pathway enzymes by separate *Agrobacterium* leads to at least some level of co-infection into the same plant cell. However, we do not know about the possibility of enzyme and/or metabolite sharing between plant cells. If these scenarios do occur at appreciable levels, this would only increase the likelihood of complete pathway reconstitution in the plant leaf. In designing *Agrobacterium*-mediated transient expression co-infiltration experiments, we show the main experimental variable determinants of co-infection rates are *Agrobacterium* infiltration OD_600_s, and leaf age (plant pavement cell size increasing with leaf age). Co-infection rates are positively correlated with both of these variables.

Beyond reconstitution applications requiring high rates of co-infiltration, applications where at most 1 T-DNA type/plant cell is desired can be reached through these methods, e.g. for the delivery of CRISPR gRNA libraries to plant cells in an intact leaf. By dialing down individual *Agrobacterium* infiltration OD_600_ to <0.02, the majority of infected plant cells are infected with only one T-DNA type. However, there is a tradeoff with fraction of infected cells, with only ~30% of plant pavement cells being infected at 0.02 OD_600_. This limits the library size that can be delivered into leaf tissue, or requires robust selection/screening methods to pull out successful infection events.

We anticipate that measurements of MOI for the *Agrobacterium*/*N. benthamiana* system will help with the design of future work requiring delivery of multiple unique T-DNA constructs and/or achieving at most 1 unique construct per cell. Furthermore, our investigation sheds light on the ease at which *N. benthamiana* can undergo transient infection with this *Agrobacterium* system, and hopefully will help elucidate why equivalent DNA-delivery methods in e.g. monocots are more challenging.

## Experimental

### Fluorescent protein cloning into pEAQ-HT vector

Fluorescent proteins were cloned into the pEAQ-HT vector ^6^ by standard cloning techniques, using the built in AgeI/XhoI cloning site. Primers 5’-TGAGAT**ACCGGT**<u>ATGGTGAGCAAGGGCGAG</u> (forward primer, added AgeI site in bold) and 5’-TGAGAT**CTCGAG**<u>TTAAGATCTGTACAGCTCGTC</u> (reverse primer, added XhoI site in bold) were used to amplify each fluorescent protein with polymerase chain reaction (PCR) with Phusion polymerase (NEB). Templates for PCRs were pAN579 (CFP), pAN581 (YFP) & pAN583 (mCherry), plasmids courtesy of the Nebenführ lab. Column purified (Zymo) PCR products and purified plasmid pEAQ-HT were each digested with AgeI-HF and XhoI (NEB) for 1 hour at 37 °C. After 30 minutes, CIP (NEB) was added to the pEAQ-HT digestion to dephosphorylate the backbone. Digestion products were column purified (Zymo), and eluted with nuclease-free water. Three ligation reactions were set up with T4 Ligase (NEB), 50 ng pEAQ-HT backbone, and 5-fold molar excess of fluorescent protein insert. Reactions were left at room temperature for 10 minutes and deactivated at 65 °C for 10 minutes, and 5 μL each were transformed into chemically competent *E. coli* DH5ɑ (NEB). Following recovery in SOC at 37 °C for 1 hour, cells were plated on LB-agar supplemented with 30 μg mL^−1^ kanamycin, and incubated overnight at 37 °C. Colonies were picked into 5 ml LB supplemented with 30 μg ml^−1^ kanamycin, grown overnight at 37 °C, and miniprepped (Zymo) for sequencing.

### Preparation of competent GV3101/pMP90

*Agrobacterium tumefaciens* GV3101/pMP90 is a C58 lineage strain, with the T-DNA portion of the wild-type virulence plasmid pTiC58 replaced with a gentamycin resistance cassette to create pMP90 ^5^. GV3101/pMP90 is grown at 30 °C, and is resistant to gentamycin (15 μg mL^−1^) from pMP90, and rifamycin (15 μg mL^−1^) from the GV3101 genome. GV3101/pMP90 is grown at 30 °C, and can be grown in standard LB.

To prepare chemically competent GV3101/pMP90, −80 °C stock was streaked out on LB-Agar plates supplemented with 15 μg mL^−1^ gentamycin and incubated at 30 °C for 2 days until clear colonies formed. A colony was picked to inoculate 5 mL of LB supplemented with 15 μg mL^−1^ gentamycin and grown to saturation at 30 °C in a roller drum. The saturated culture was added to 1 L of LB supplemented with 15 μg mL^−1^ gentamycin in a baffled flask and incubated at 30 °C with 250 rpm shaking until an OD_600_ of ~0.5 was reached. The culture flask was transferred to ice, shaken vigorously to rapidly cool the culture, and left on ice for 20 minutes with occasional shaking. Culture was pelleted in a pre-cooled (4 °C) centrifuge for 5 min at 4,000 xg. Supernatant was removed, and the pellet resuspended in 20 mL ice cold 20 mM CaCl_2_. The cell suspension was aliquoted (100 μL each) into 1.5 mL Eppendorf tubes, flash frozen in liquid nitrogen, and stored at −80 °C until use.

### Transformation of competent GV3101/pMP90

Purified pEAQ-HT plasmid DNA was added to an aliquot of frozen competent GV3101/pMP90 cells. Because of the low transformation efficiencies, ~1 μg of DNA should be used (in this work, 5 μL was added of ~200 ng μL^−1^ plasmid stock). The Eppendorf tube was placed in a 37 °C heat block for 5 minutes, and then mixed well by flicking the tube 5-10 times. The cells were flash frozen on liquid nitrogen and thawed at 37 °C for 5 minutes. 1 mL of LB was added, and the tube incubated at 30 °C with rotation for 2 hours. Cells were pelleted in a microcentrifuge for 4 minutes at 5,000 xg. All but ~100 μL of supernatant was removed and cells resuspended. Concentrated cells were spread with sterile ~4 mm glass beads on LB-agar plates supplemented with 15 μg mL^−1^ gentamycin and 30 μg mL^−1^ kanamycin, and incubated at 30 °C for 2 days until colonies appeared. A colony was then re-streaked onto LB-agar agar plates supplemented with 15 μg mL^−1^ gentamycin and 30 μg mL^−1^ kanamycin, and incubated at 30 °C for 1-2 days until colonies appeared.

### N. benthamiana growth

*Nicotiana benthamiana* plants were grown indoors at room temperature under 16/8 hour light/dark cycle. Plants were watered twice a week with 2 g L^−1^ fertilizer (Peters Excel 15-5-15).

### Growth, induction, and infiltration of GV3101 strains into N. benthamiana leaves

A colony of GV3101/pMP90/pEAQ-[CFP/YFP/mCherry] was picked with a sterile P10 pipette tip into 5 mL LB supplemented with 15 μg mL^−1^ gentamicin and 30 μg mL^−1^ kanamycin, and incubated for 24 hours at 30 °C with shaking (250 rpm). Cell culture was transferred to a 15 mL falcon tube, pelleted at 5000 xg for 5 minutes, and resuspended in 4 mL of induction buffer (10 mM MES buffer pH 5.6; 10 mM MgCl_2_; 150 μM acetosyringone). Cell suspensions were left at room temperature for 1-2 hours, occasionally inverting to mix. Following induction, the OD_600_ of the culture was taken (Thermo Scientific Genesys 20), and culture OD_600_ was normalized to 0.6 OD_600_ with fresh induction buffer. Infiltration mixes were prepared with these induced cultures, depending on the experiment. *Agrobacterium* mixes were hand infiltrated into the underside of the leaf of 6-week old *Nicotiana benthamiana* using a 1 mL blunt-end syringe. Infiltrated plants were returned to 18/6 light/dark cycle growth shelf for 5 days.

### Confocal microscopy imaging

Vacuum grease was applied near the edge of a square coverslip (VWR 48366-223) with a 3 mL Luer lock syringe fitted with a blunt-end 18G needle, enough to form a continuous seal of ~1 mm. Water (~20 μL) was pipetted onto the center of the coverslip. The leaf to be imaged was cut from the plant, and a leaf disk punched (#4 size) from an infiltrated section of the leaf, and placed underside (abaxial) side down on the water spot. A microscope slide (Fischer Scientific microscope slide 12-500-A3) was placed on top, and pressed evenly and gently to form a seal with the coverslip and vacuum grease. A Leica TCS SP8 laser scanning confocal microscope in resonant scanning mode with LASX software was used to visualize fluorescent protein expression in leaves with a 20x dry objective. CFP fluorescence was imaged with excitation 440 nm/emission 470/15 nm, GFP fluorescence with excitation 488 nm/emission 525/50 nm, YFP fluorescence with excitation 515 nm/emission 550/30 nm, and RFP or mCherry fluorescence with excitation 580 nm/emission 610/20 nm. Microscope images were processed using Fiji-ImageJ. Projection type of Max Intensity was used to flatten z-stacks, and Enhance Contrast with 0.3% saturated pixels and equalize histogram was used (except for Figure 2 image processing, which were not contrast enhanced because of pixel intensity level quantification).

### Quantifying plant pavement cell perimeters

Microscope images were processed and printed, then plant pavement cells fully within frame were traced onto blank paper by hand with a backlit LED light box. Traced images were scanned and transformed into a vector image using Adobe Illustrator pathfinder function (Supplementary figures 9 and 10A). Individual plant pavement cell perimeter was calculated using Adobe Illustrator; Window>Document Info, then select “Object” from top right pull-down menu in the new window, then select an object, and path length will be displayed. Co-infection events were manually counted using ImageJ, counting a cell positive for a given fluorescent protein if the key features of cell perimeter and nucleus could be clearly observed in that channel and cell.

### Poisson Distribution modeling equations

To model infection of plant cells by *Agrobacterium*, we assume that an *Agrobacterium* cell infiltrated into a leaf can lead to productive infection (*i.e*., infection that leads to gene expression) with some unknown probability, and that infection events are mutually independent. If these conditions are satisfied, the number of productive *Agrobacterium* infection events per infectable plant cell (*X*) can be modeled using a Poisson distribution:

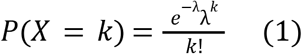

Where *k* is a non-negative integer and λ is the average number of infection events per cell. Various factors such as leaf age, plant cell size, or the nature of the T-DNA may affect λ, but under otherwise identical experimental conditions, λ will be directly proportional to the total number of *Agrobacterium* cells infiltrated, which can be quantified via an OD_600_ measurement.

We therefore have:

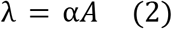

where *A* denotes the OD_600_ and α is a scaling factor. The probability that a cell is not infected (which we also denote as *f*_0_) is then

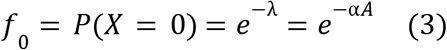

while the probability that a cell is infected at least once is therefore

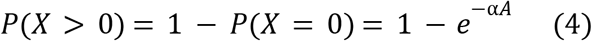

To determine α, we can vary *A* for an indicator strain carrying a fluorescent protein expression T-DNA and measure the fraction of plant cells that do not express this protein, *i.e*. *f* _0_. The best fit value for α can be determined from experimental data by linear regression using a suitably transformed version of eqn. 3:

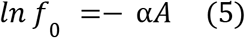

To model co-infection by multiple *Agrobacterium* strains, we assume that α is independent of the gene being expressed. Since infection events are mutually independent, we have

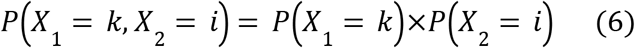

for any pair of strains. Hence, for *n* different strains, the probability that a cell is infected by every strain at least once is given by:

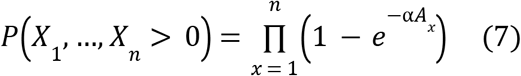

Where *A*_*x*_ is the OD_600_ of strain *x*. If each strain carries a biosynthetic pathway enzyme, for example, eqn. 7 gives the fraction of cells expected to express the entire pathway. For the case 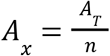, where the OD_600_ of each infiltrated strain is the same, eqn. 7 simplifies to

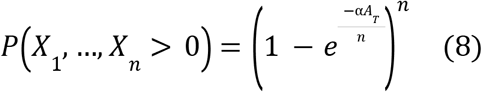

Finally, for a pair of strains with *A* 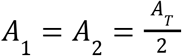, we have:

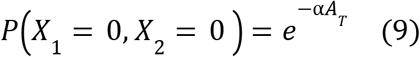

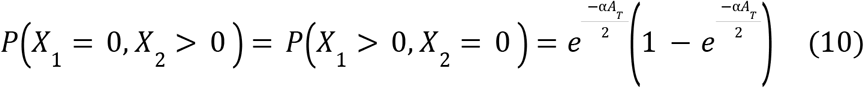

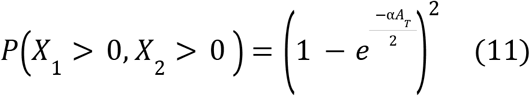

Eqns. 10 and 11 can be used to estimate α in a dataset where α*A*_T_ is such that 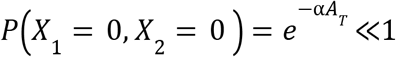, but *P*(*X*_1_ > 0, *X* _2_ > 0) is reasonably small (between 0.1 and 0.8, say).

## Acknowledgements

The authors would like to thank Heather Cartwright and Andrey Malkovskiy of the Carnegie Science Department of Plant Biology for training and use of the Leica SP8 Confocal Microscope, and Will B. Cody for manuscript comments.

## Data Availability Statement

All data is included in the supplementary information and data file, and microscopy data available in the Image Data Resource.

---

**Supplementary Figure 1.**
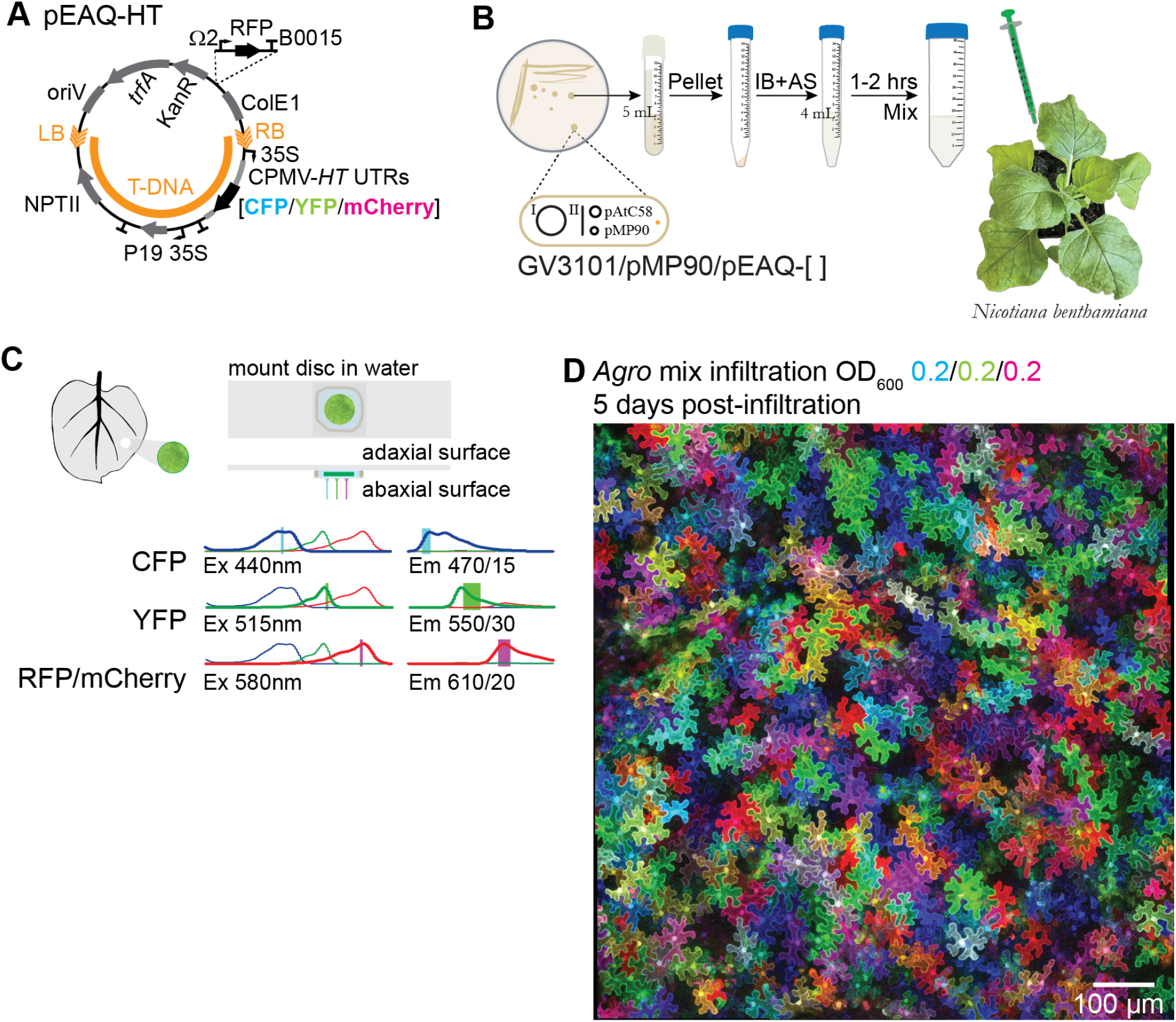
*Agrobacterium*-mediated transformation for transient heterologous protein expression and confocal microscopy imaging. **A**. pEAQ-HT expression vector map. **B**. *Agrobacterium*-mediated transient expression infiltration method. **C**. Confocal microscopy imaging settings. **D**. Confocal microscope image of underside of leaf, 5 days post-infiltration, 0.2 OD_600_ per strain CFP/YFP/mCherry, 0.6 OD_600_ total.

**Supplementary Figure 2.**
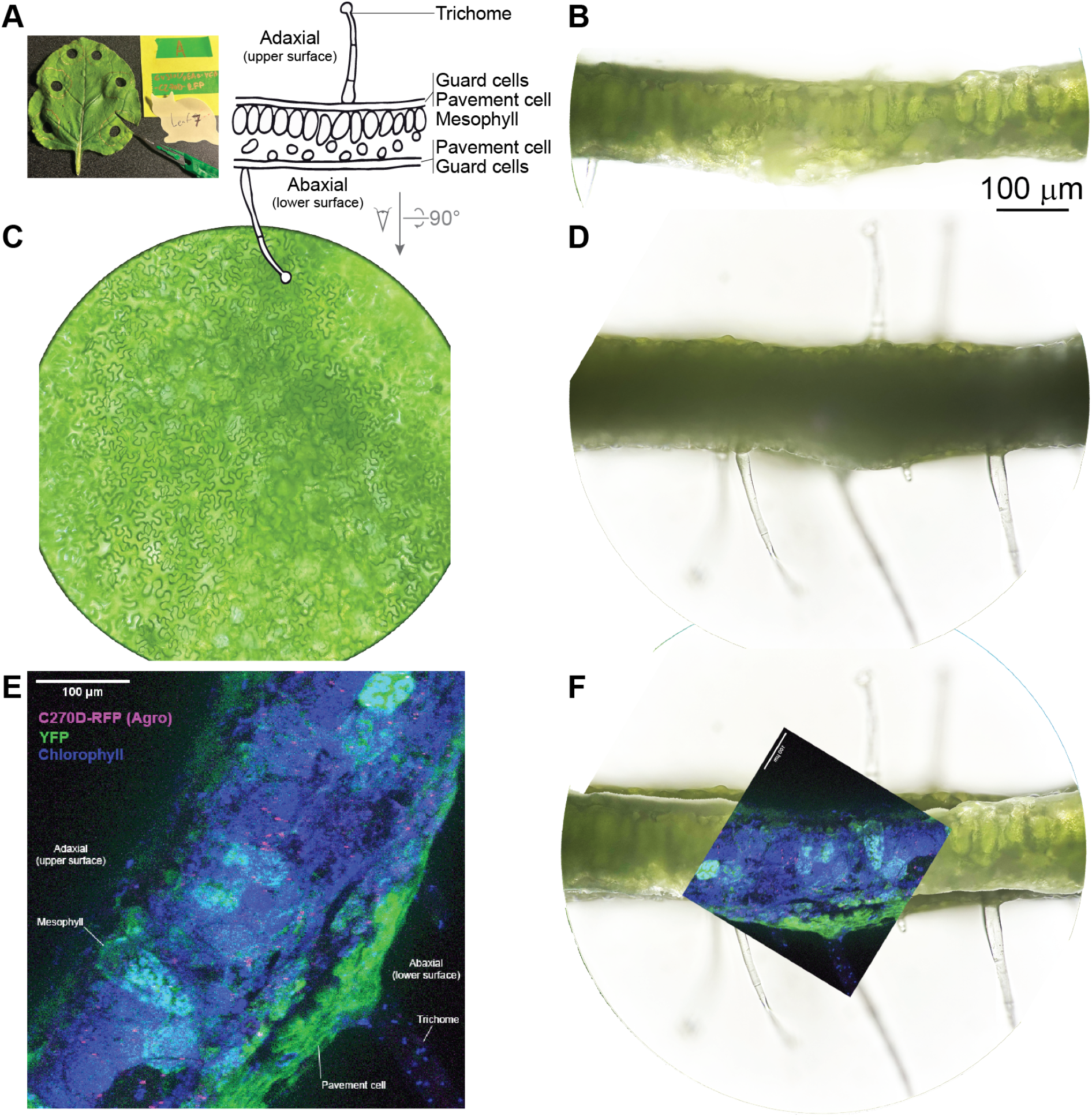
Brightfield and confocal microscopy images used to construct *N. benthamiana* leaf cross section diagram. **A**. Thin cross section cut from a 5 dpi infiltration spot, infiltrated with GV3101(pMP90)/pEAQ-YFP-RFP, with fresh scalpel. **B**. Brightfield image of leaf cross-section, focused on cut surface. **C**. Bright field image of Abaxial (lower) surface of leaf. **D**. Bright field image of leaf cross-section, focused further in from cut surface. **E**. Confocal microscopy image of RFP in magenta (*Agrobacterium*), YFP T-DNA product in green, and chlorophyl fluorescence in blue. **F**. Overlays of images in panels **B**, **D** and **E**.

**Supplementary Figure 3.**
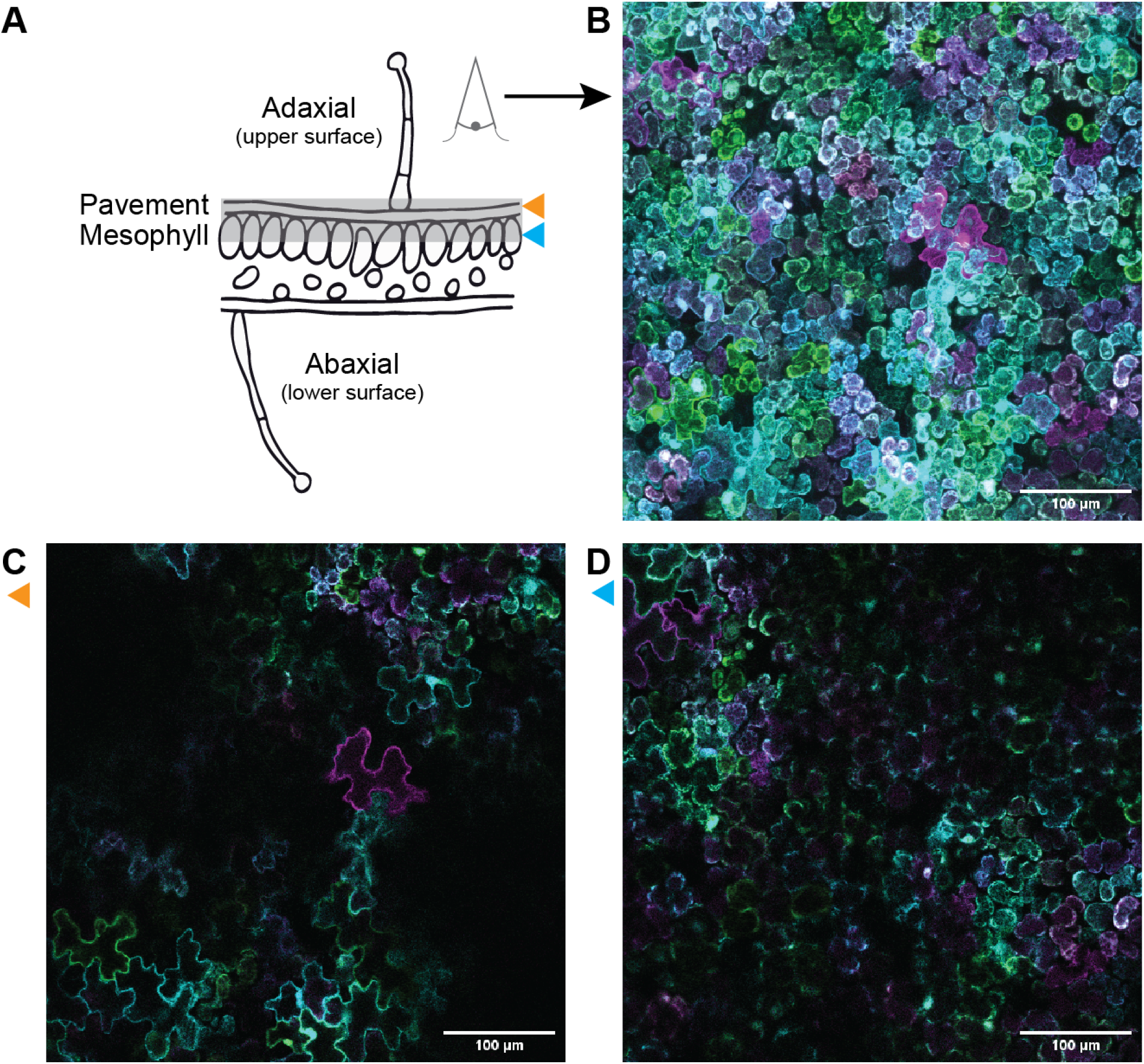
Confocal microscopy of upper surface showing transient expression in mesophyll cells. **A**. Cross section diagram of leaf. Gray box indicates full z stack projection in (**B**). Orange arrow indicates z slice across pavement cell layer (**C**). Blue arrow indicates z slice across mesophyll cell layer (**D**). White bar indicates 100 μm.

**Supplementary Figure 4.**
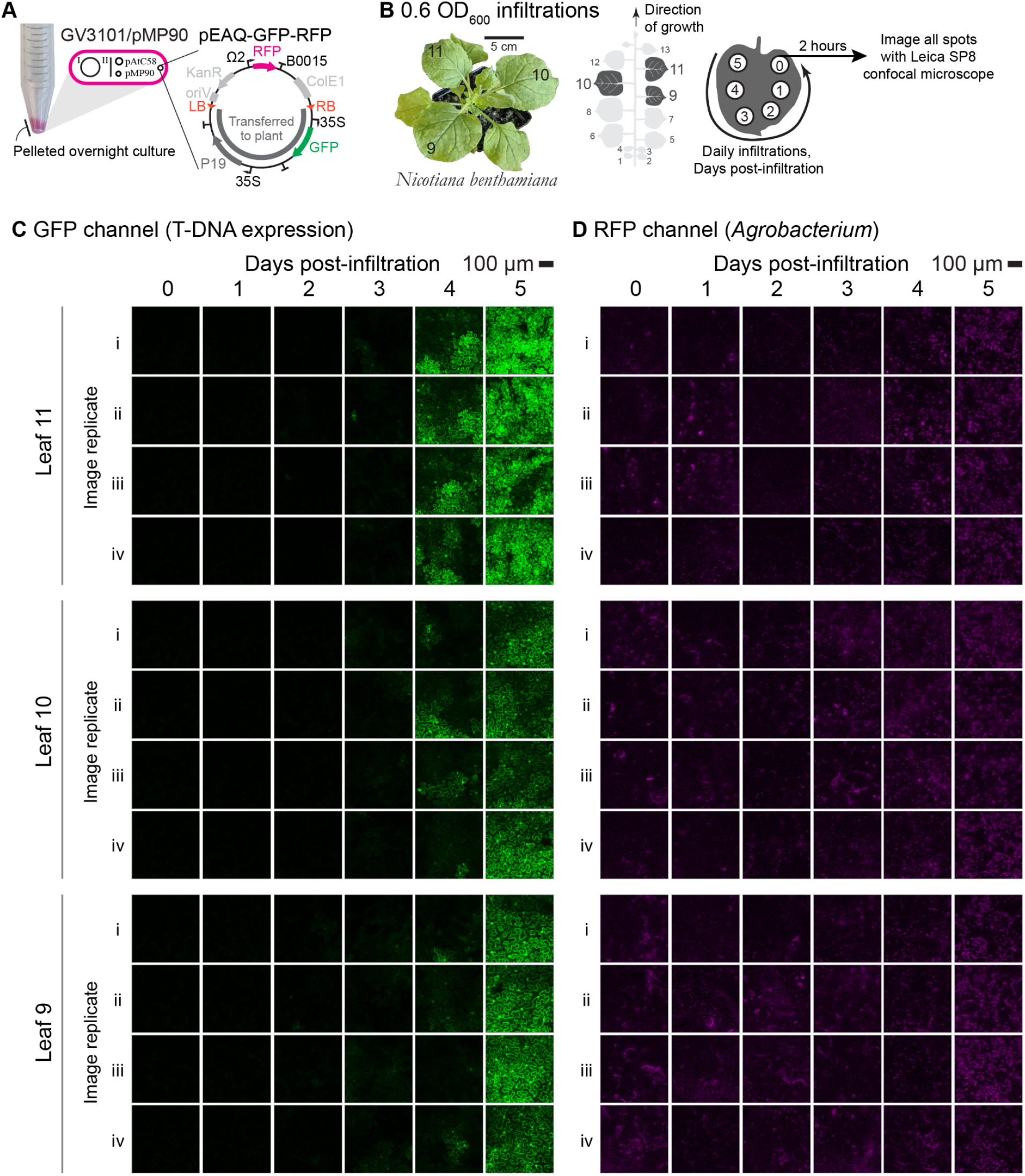
Tracking of RFP-labeled GV3101 strain with a GFP T-DNA load at 0.6 infiltration OD_600_. **A**. GV3101/pMP90/pEAQ-GFP-RFP strain. **B**. Photo of *Nicotiana benthamiana* 6-week old plant and leaf numbering schematic, with the oldest cotyledons counted as 1 and 2, increasing to the youngest leaf. **C**. GFP channel. **D**. RFP channel.

**Supplementary Figure 5.**
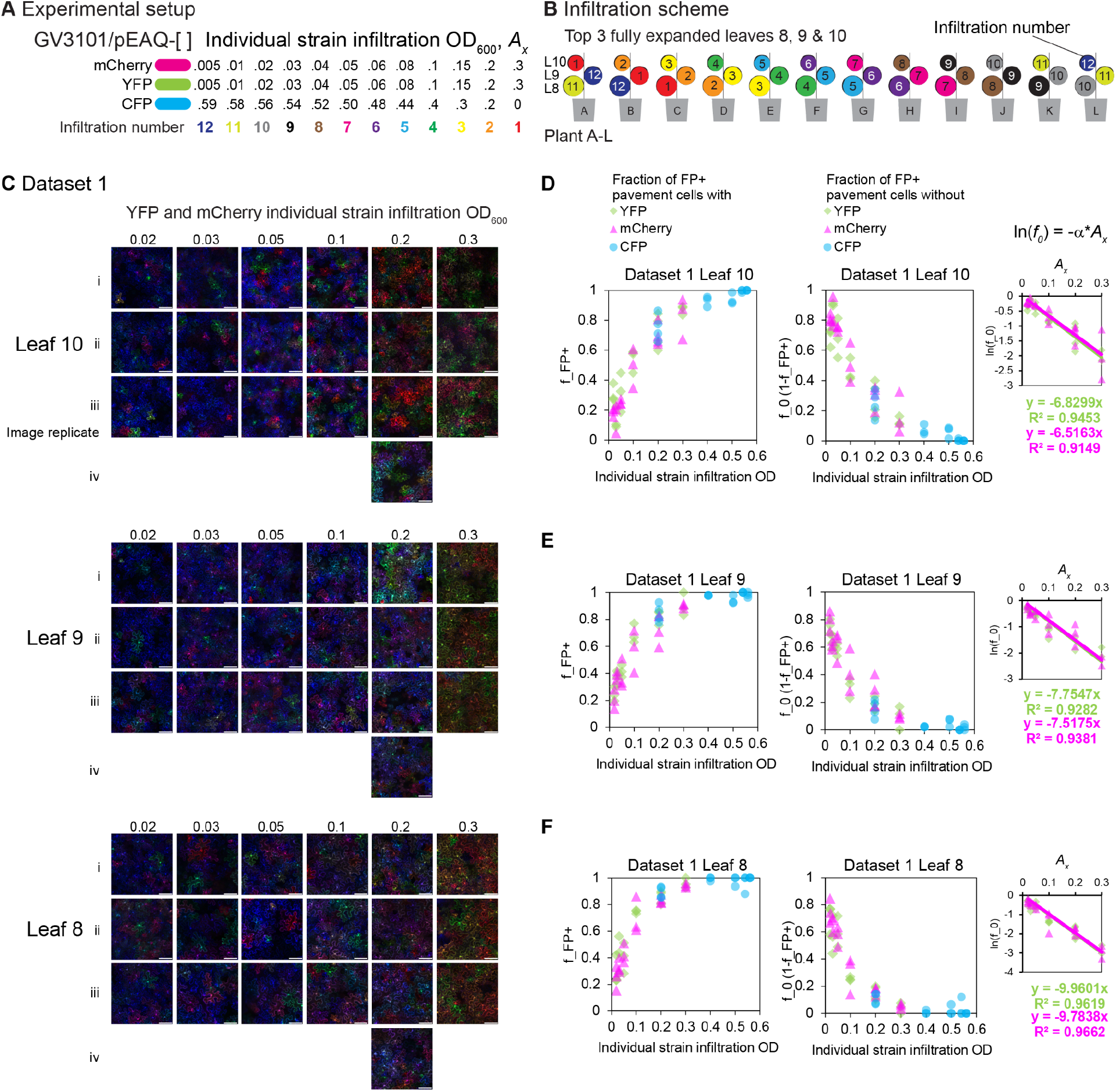
Multiplicity of infection determination, Dataset 1. **A,B**. Experimental setup. **A**. *Agrobacterium* strain co-infiltration mixes **B**. Infiltration scheme into the top 3 fully expanded leaves. **C**. Overview of microscopy image data collected for Dataset 1. **D-F**. Determining α values for Leaf 10 (**D**), Leaf 9 (**E**) and Leaf 8 (**F**).

**Supplementary Figure 6.**
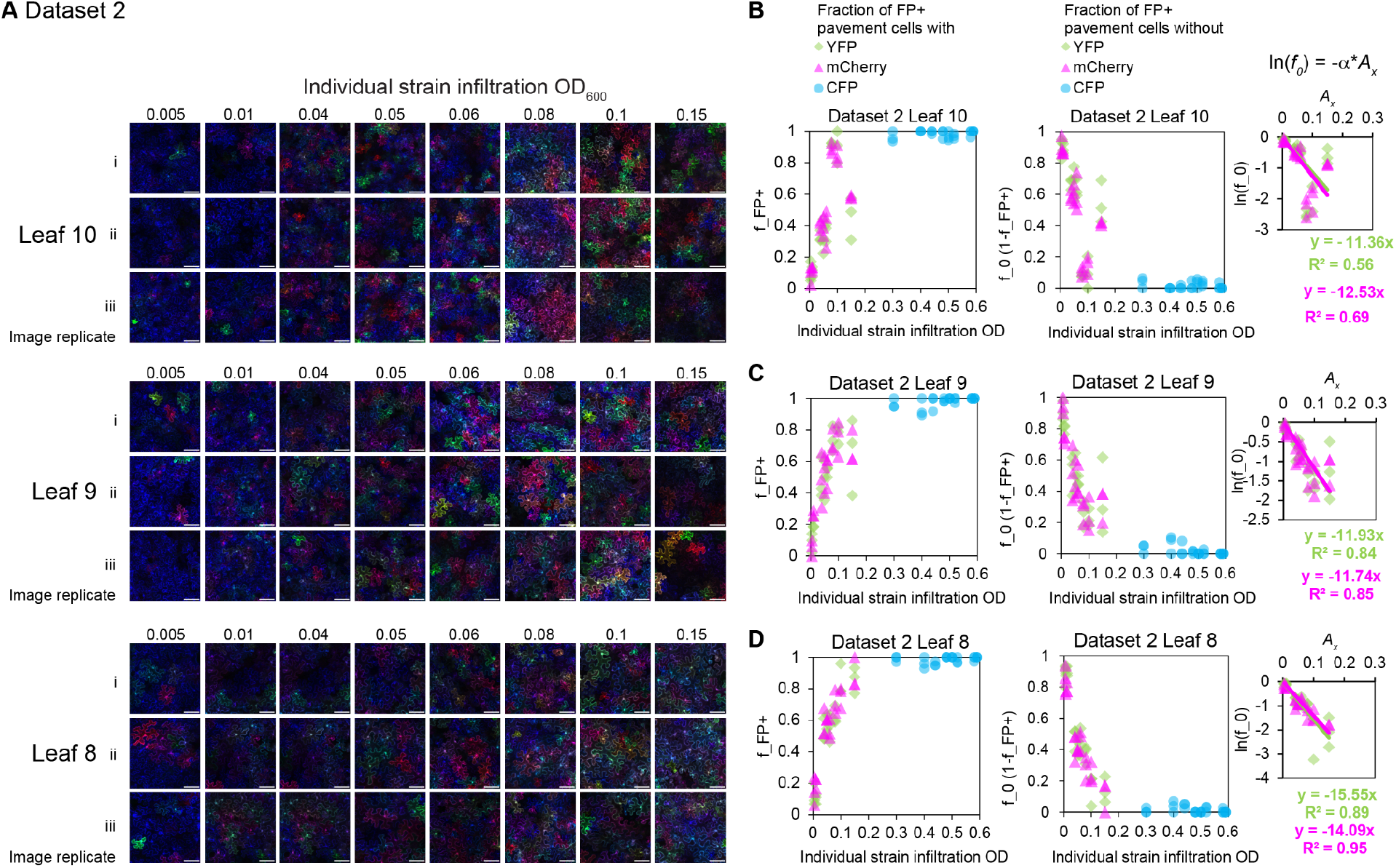
Multiplicity of infection determination, Dataset 2. **A**. Overview of microscopy image data collected for Dataset 1. **B-D**. Determining α values for Leaf 10 (**B**), Leaf 9 (**C**) and Leaf 8 (**D**).

**Supplementary Figure 7.**
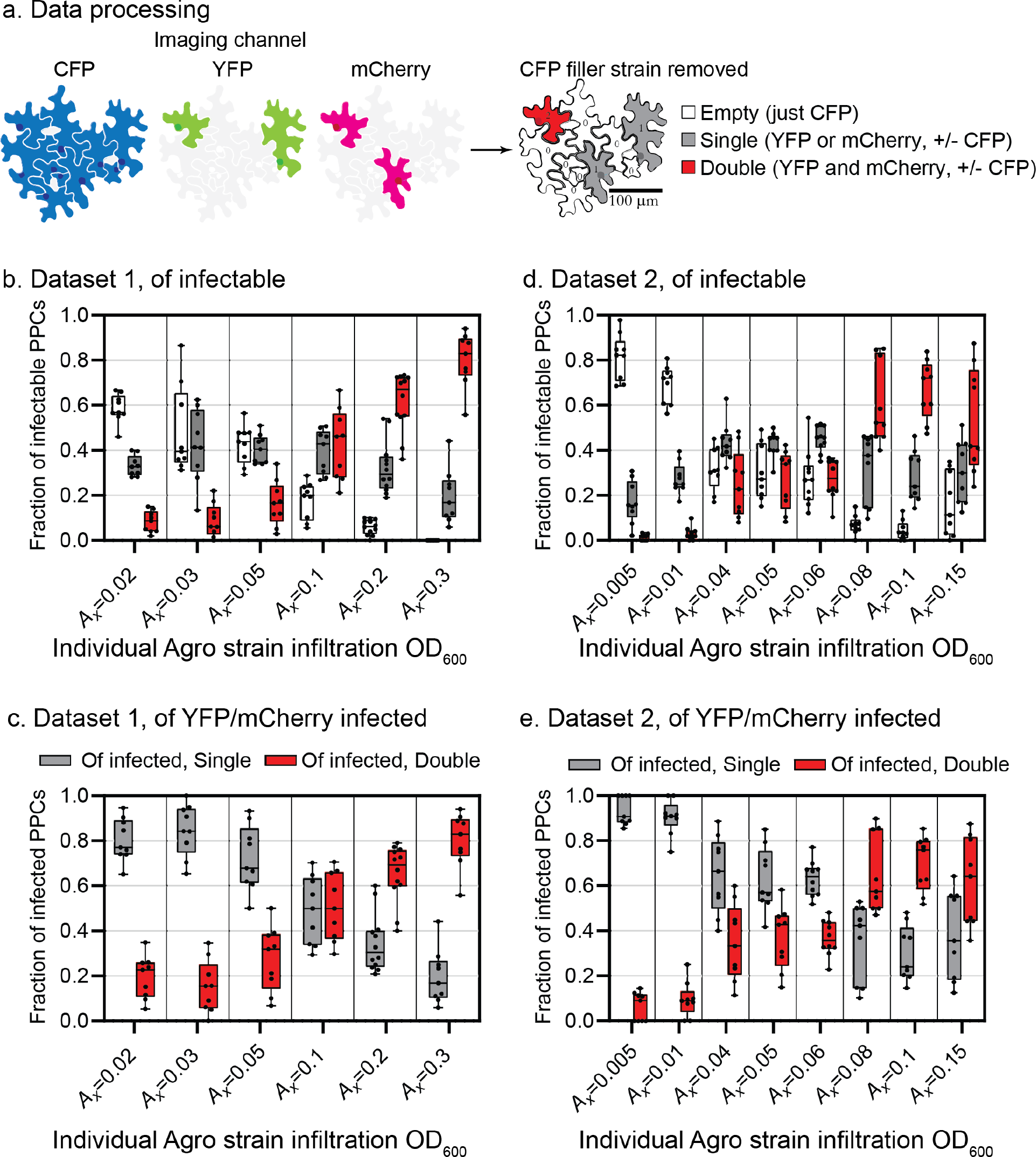
**A**. Experimental data processed for YFP and mCherry indicator strains, empty, single and double infection bins. **B, D**. Fraction of infectable plant pavement cells (PPCs) with no YFP or mCherry (empty), YFP or mCherry (single), and YFP and mCherry (double) for Dataset 1 (**B**) and Dataset 2 (**D**). **C, E**. Data processed for only indicator strain positive pavement cells (YFP and/or mCherry) for Dataset 1 (**C**) and Dataset 2 (**E**).

**Supplementary Figure 8.**
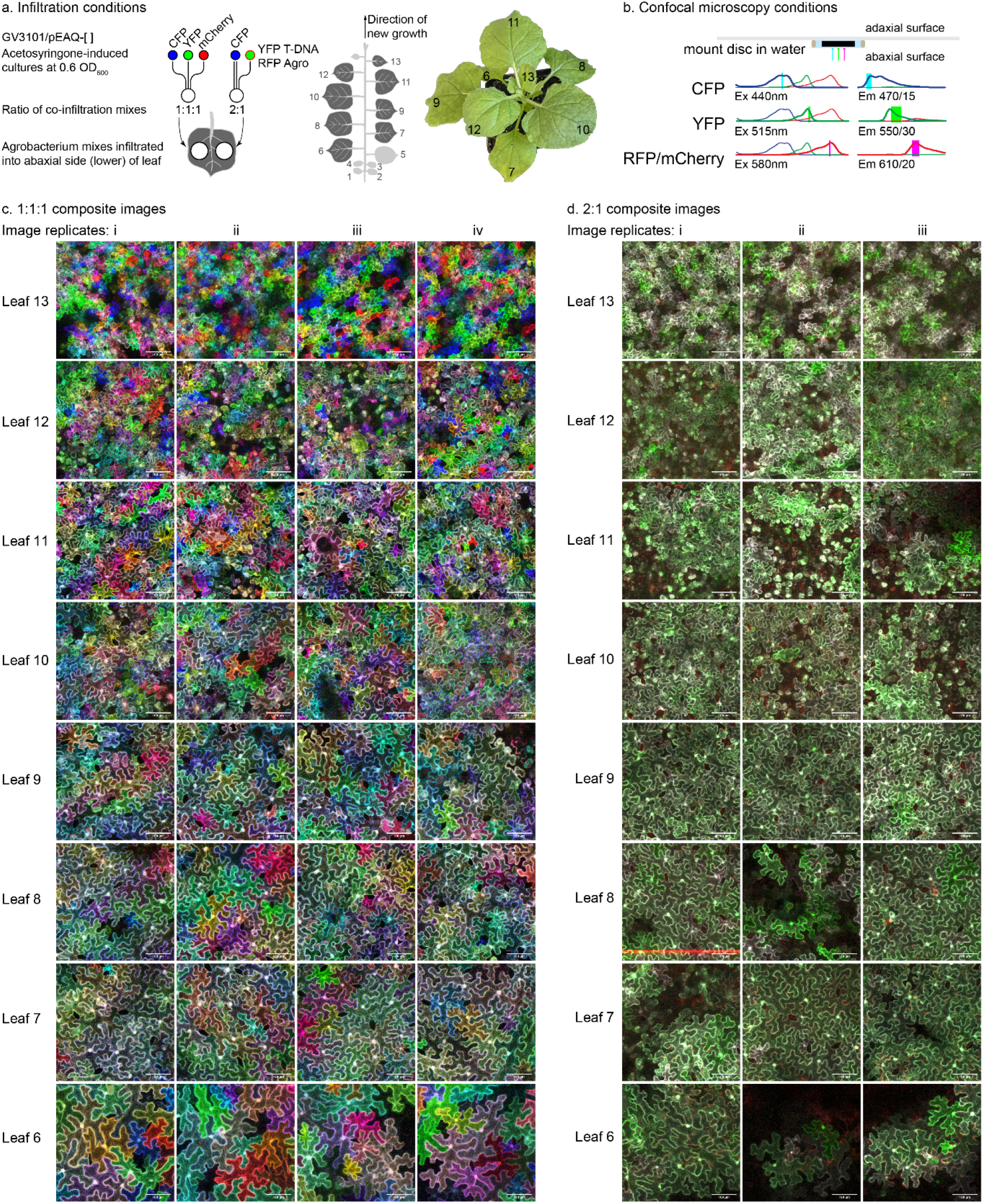
Microscopy imaging dataset for Figure 4. **A**. Infiltration scheme of Acetosyringone-induced culture mixes. On left side of leaf, 1:1:1 mix of CFP:YFP:mCherry cultures, each at a final OD_600_ of 0.2, total infiltration OD_600_ 0.6. On right side of leaf, a 2:1 mix of CFP strain (0.4 OD_600_) with a YFP (T-DNA) and RFP (*Agro* tag) strain (0.2 OD_600_). Mixes were infiltrated into leaves 6-13 on a 6-week old *N. benthamiana* plant. **B**. Confocal microscopy imaging conditions. **C**. Microscopy dataset for 1:1:1 CFP:YFP:mCherry infiltration spot, 4 image replicates per leaf. Composite image shown with CFP channel colored blue, YFP channel colored green, and mCherry channel colored red. White scale bar indicates 100 μm. **D**. Microscopy dataset for 2:1 CFP:YFP-RFP infiltration spot.

**Supplementary Figure 9.**
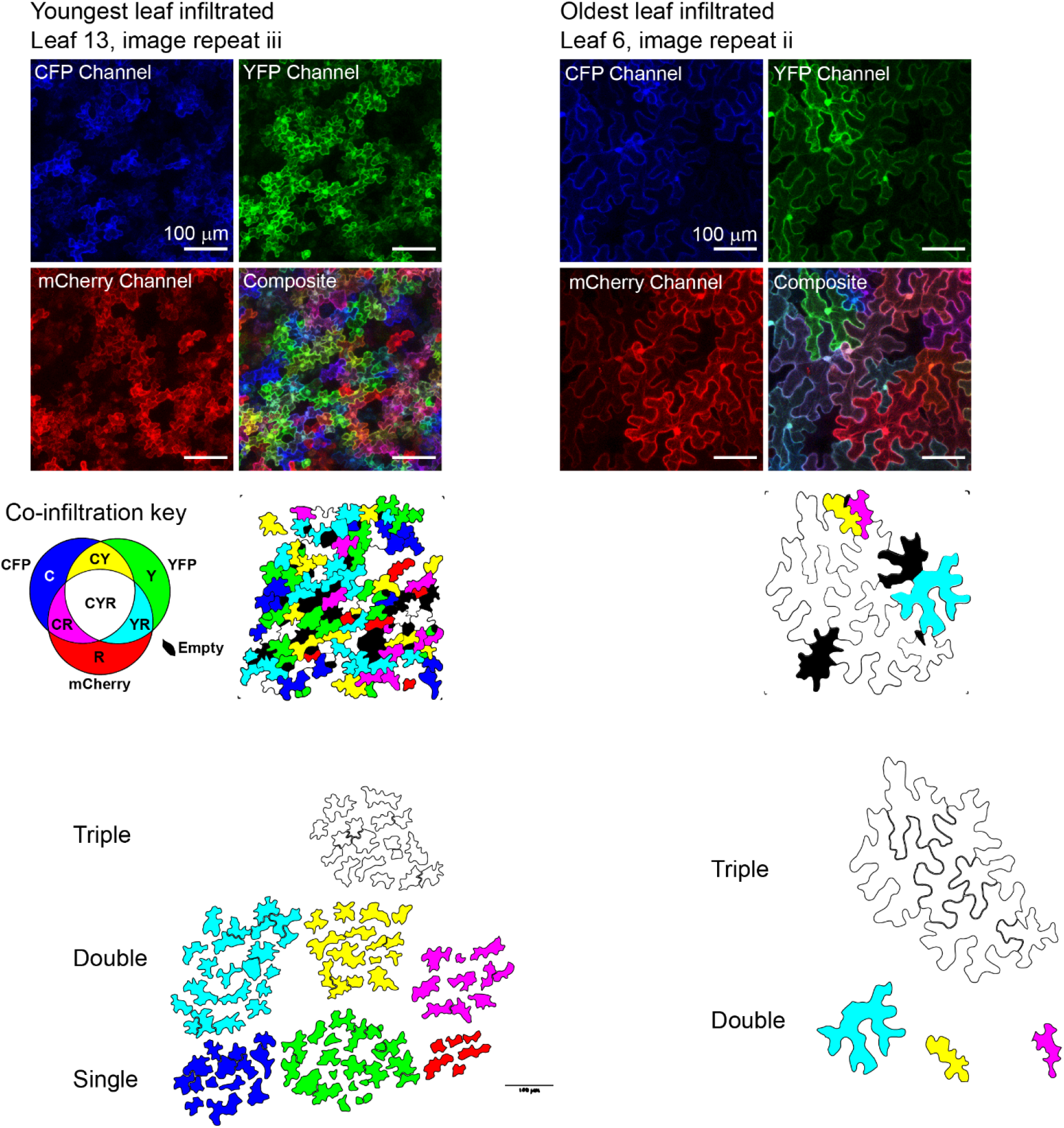
Image processing and quantification procedure and example. Representative traced and counted images. Co-expression key: C: CFP, Y: YFP, R: mCherry. A white plant pavement cell indicates co-infection by all 3 strains, black, no detectable fluorescent protein. White scale bar 100 μm.

**Supplementary Figure 10.**
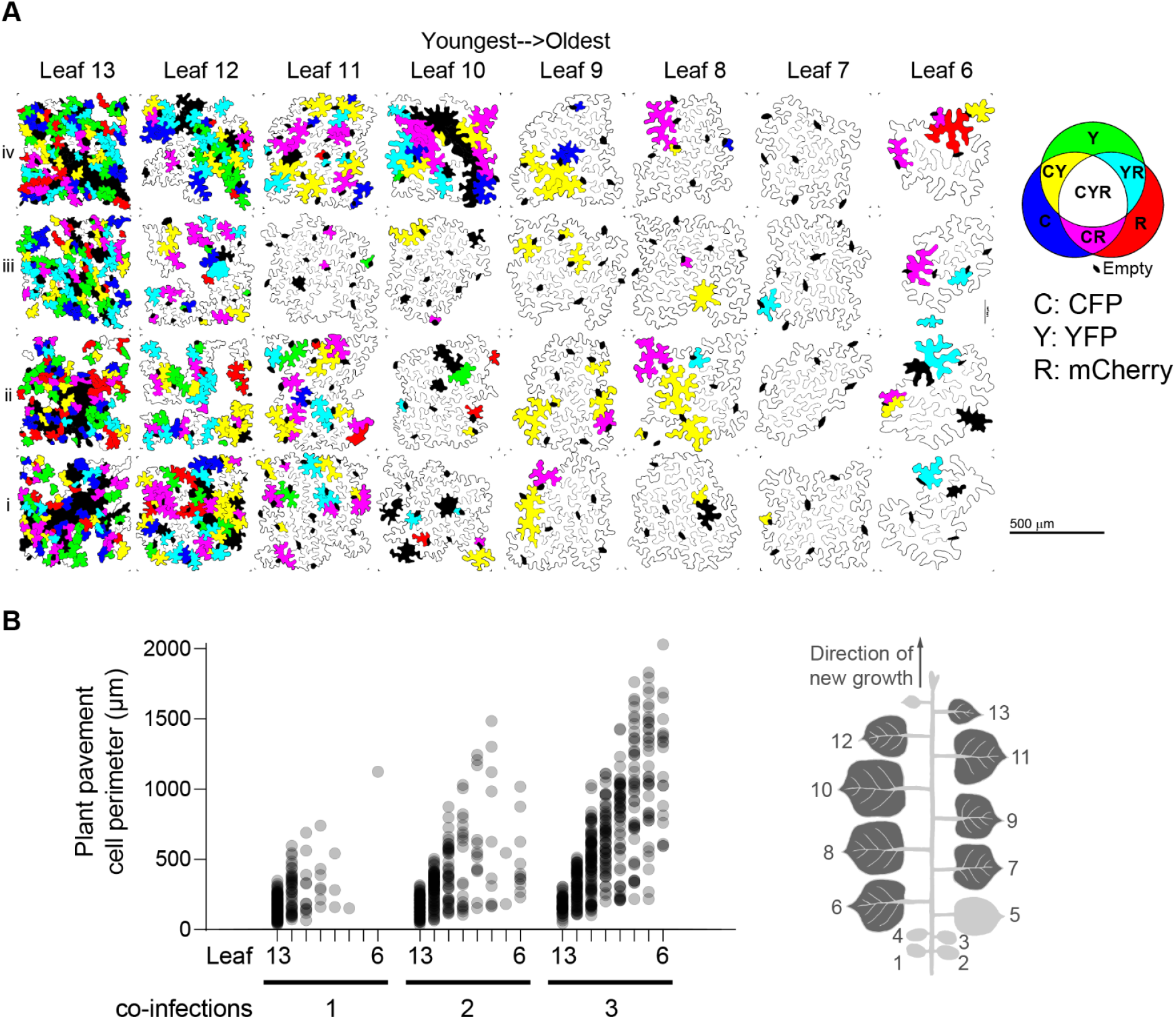
Analysis of data from 1:1:1 CFP:YFP:mCherry strain co-infiltration, 0.2 OD_600_ per strain, total infiltration OD_600_ = 0.6. **A**. Plant pavement cell traces for perimeter quantification and co-infection binning. C: CFP, Y: YFP, R: RFP. **B**. Co-infections and pavement cell perimeters by leaf.

**Supplementary Figure 11.**
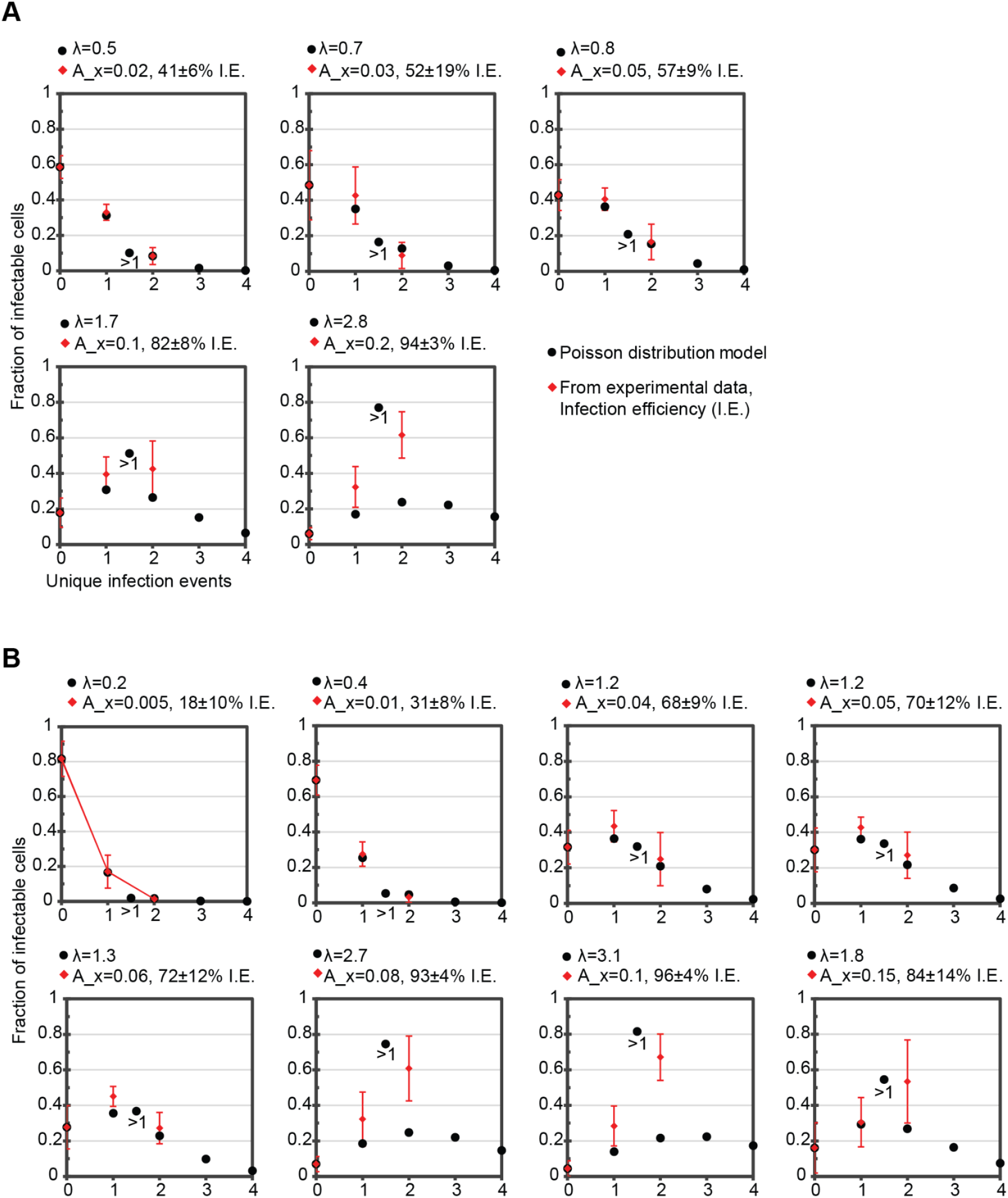
Expected number of infection events per plant pavement cell at various infection efficiencies based on Poisson distribution (equation 1). Curves with solid connection lines are based on the Poisson distribution form, with overlaid comparison of experimental data in red square with dashed line. Datasets 1 (**A**) and 2 (**B**).

**Supplemental Table 1.**
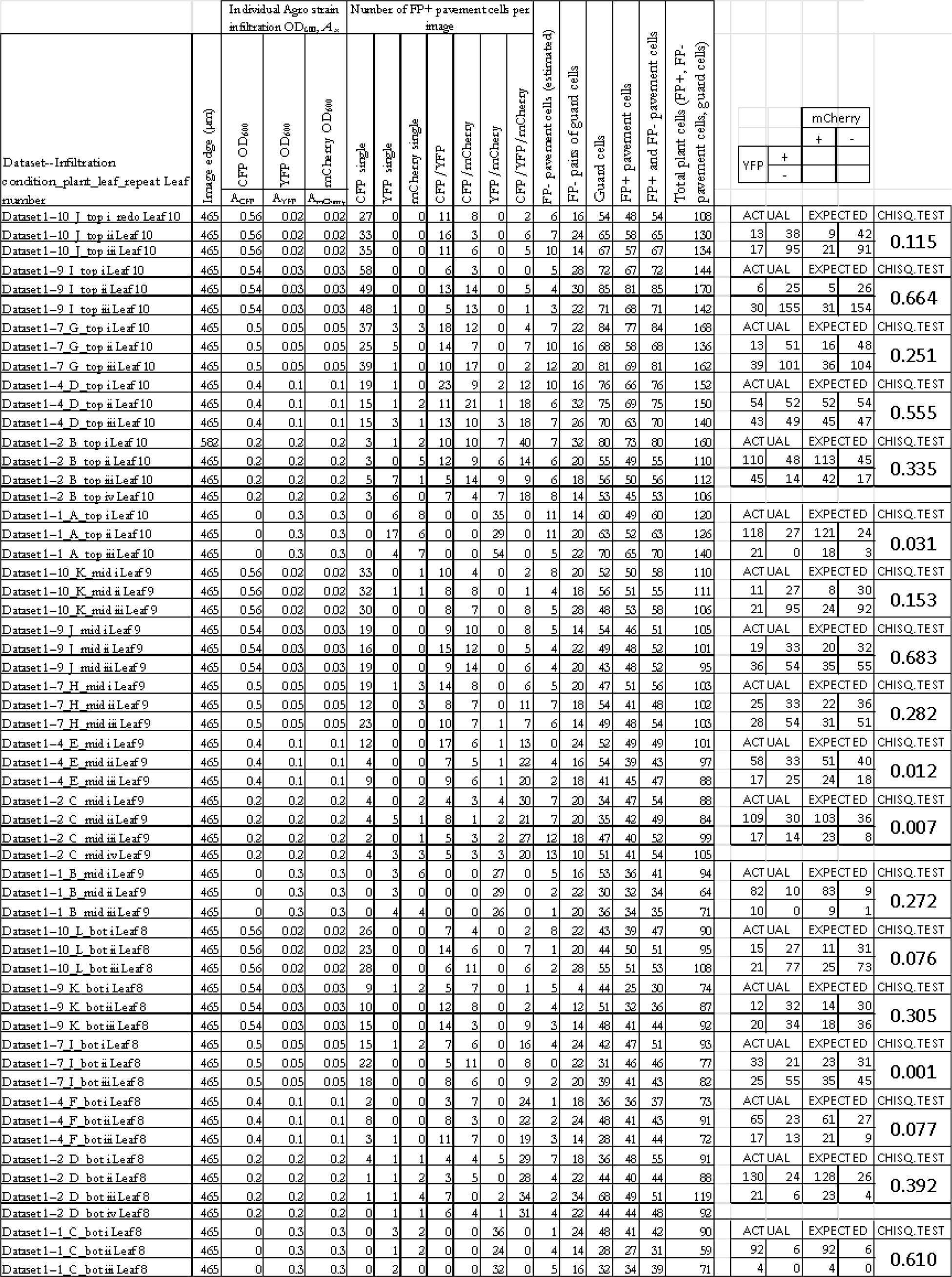
Data for figure 3, Dataset 1.

**Supplemental Table 2.**
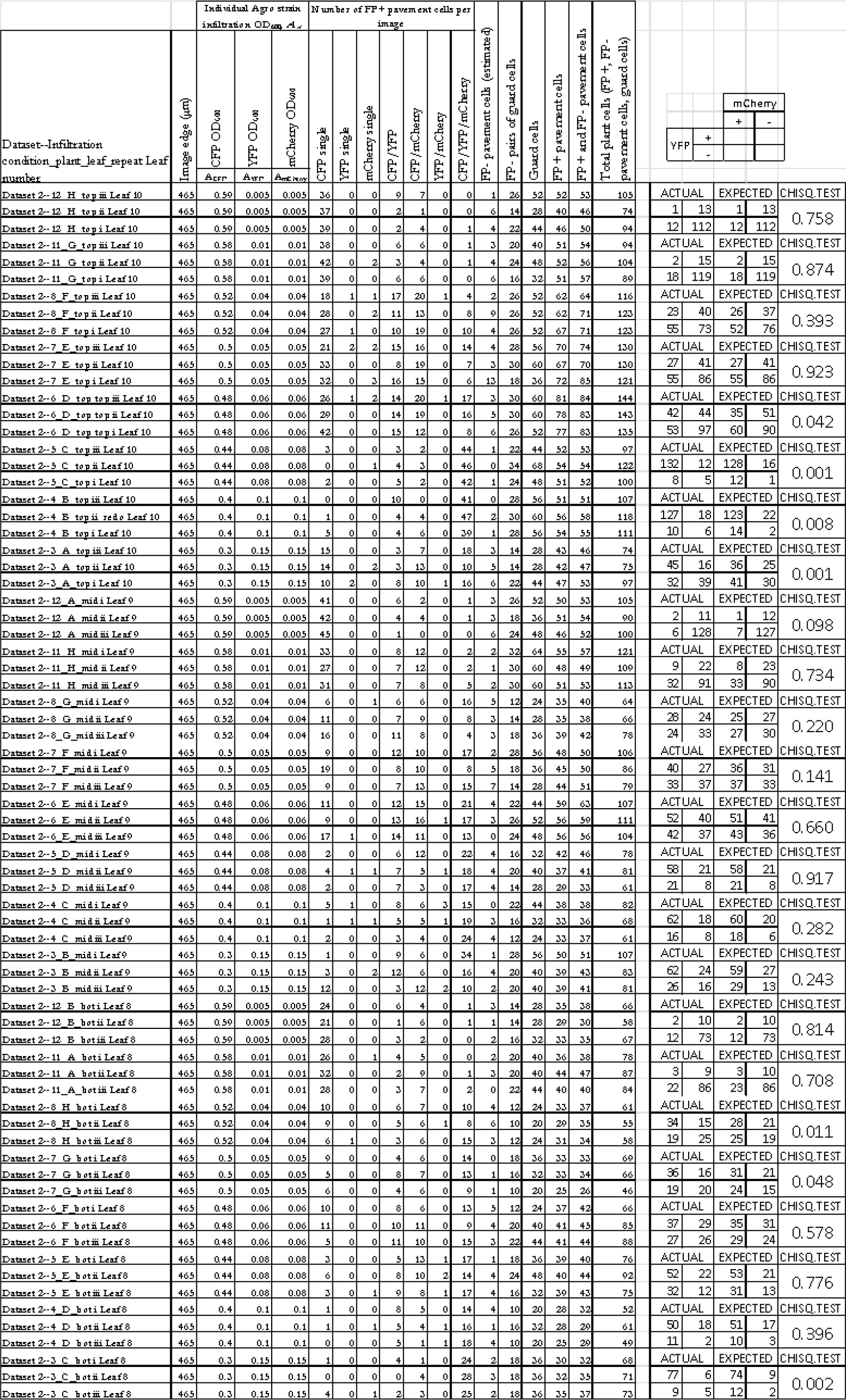
Data for figure 3, Dataset 2

**Supplemental Table 3.** Data for figure 3

| Leaf\Image | PPCs with 1 infection, n_1 |  |  |  | PPCs with 2 infections, n_2 |  |  |  | PPCs with 3 infections, n_3 |  |  |  | Total infected PPCs |  |  |  | f_1, Fraction of infected w/1 |  |  |  | f_2, Fraction of infected w/2 |  |  |  | f_3, Fraction of infected w/3 |  |  |  |
| --- | --- | --- | --- | --- | --- | --- | --- | --- | --- | --- | --- | --- | --- | --- | --- | --- | --- | --- | --- | --- | --- | --- | --- | --- | --- | --- | --- | --- |
|  | i | ii | iii | iv | i | ii | iii | iv | i | ii | iii | iv | i | ii | iii | iv | i | ii | iii | iv | i | ii | iii | iv | i | ii | iii | iv |
| 13 | 42 | 54 | 42 | 57 | 39 | 30 | 46 | 39 | 19 | 7 | 17 | 11 | 100 | 91 | 105 | 107 | 0.4 | 0.6 | 0.4 | 0.5 | 0.4 | 0.3 | 0.4 | 0.4 | 0.2 | 0.1 | 0.2 | 0.1 |
| 12 | 13 | 10 | 6 | 15 | 28 | 27 | 17 | 18 | 17 | 16 | 27 | 15 | 58 | 53 | 50 | 48 | 0.2 | 0.2 | 0.1 | 0.3 | 0.5 | 0.5 | 0.3 | 0.4 | 0.3 | 0.3 | 0.5 | 0.3 |
| 11 | 1 | 4 | 1 | 6 | 13 | 13 | 2 | 16 | 31 | 20 | 24 | 22 | 45 | 37 | 27 | 44 | 0 | 0.1 | 0 | 0.1 | 0.3 | 0.4 | 0.1 | 0.4 | 0.7 | 0.5 | 0.9 | 0.5 |
| 10 | 1 | 3 | 0 | 3 | 4 | 2 | 3 | 11 | 20 | 15 | 18 | 6 | 25 | 20 | 21 | 20 | 0 | 0.2 | 0 | 0.2 | 0.2 | 0.1 | 0.1 | 0.6 | 0.8 | 0.8 | 0.9 | 0.3 |
| 9 | 0 | 0 | 0 | 3 | 2 | 5 | 3 | 3 | 15 | 14 | 15 | 12 | 17 | 19 | 18 | 18 | 0 | 0 | 0 | 0.2 | 0.1 | 0.3 | 0.2 | 0.2 | 0.9 | 0.7 | 0.8 | 0.7 |
| 8 | 0 | 0 | 0 | 1 | 1 | 5 | 2 | 2 | 13 | 8 | 12 | 11 | 14 | 13 | 14 | 14 | 0 | 0 | 0 | 0.1 | 0.1 | 0.4 | 0.1 | 0.1 | 0.9 | 0.6 | 0.9 | 0.8 |
| 7 | 0 | 0 | 0 | 0 | 1 | 0 | 1 | 0 | 10 | 7 | 11 | 9 | 11 | 7 | 12 | 9 | 0 | 0 | 0 | 0 | 0.1 | 0 | 0.1 | 0 | 0.9 | 1 | 0.9 | 1 |
| 6 | 0 | 0 | 0 | 1 | 1 | 3 | 3 | 4 | 5 | 6 | 4 | 4 | 6 | 9 | 7 | 9 | 0 | 0 | 0 | 0.1 | 0.2 | 0.3 | 0.4 | 0.4 | 0.8 | 0.7 | 0.6 | 0.4 |

